# Functional Adaptations for Load-Bearing in a Dermal Bone: The Pectoral Fin Spine of the Russian Sturgeon (*Huso gueldenstaedtii*)

**DOI:** 10.64898/2026.04.07.716894

**Authors:** Esteban Marroquín-Arroyave, Joshua Milgram

## Abstract

Dermal bone, which forms a variety of skeletal structures and persists in a wide range of extant vertebrates, evolved prior to endochondral bone which forms all mammalian load-bearing bones. Sturgeons are a family of fish which diverged soon after the lobe-finned/ray-finned split. Sturgeon retain a long robust spine at the leading edge of the pectoral fin, called the pectoral fin spine (PFS). Pectoral fin spines are bone elements that are present in many extinct and extant species of non-tetrapod jawed fish. In this study, we characterize the structure (light, polarized, micro-computed tomography and scanning electron microscopy), composition (FTIR, TGA, BMD), and mechanical properties (3-point bending and microindentation) of the pectoral fin spine (PFS) of the Russian sturgeon (*Huso gueldenstaedtii*). The microstructure of the PFS is highly organized as it is formed by dermal osteonal bone and parallel fibered bone. Its microarchitecture, along with high material toughness, anisotropy, and substantial ash content, enables the PFS to bear loads and function in both locomotion and protection. In addition, we show an interconnected network of neurovascular canals and ornamentations, features also found in pectoral fin spines of other non-tetrapod jawed fish. Collectively, these findings demonstrate that dermal bone can form structurally organized, mechanically competent load-bearing elements and provide new insight into pectoral fin spines in ray-finned fish.

## Introduction

Bone develops either by replacement of a cartilage template or directly from mesenchymal cells without a cartilaginous model forming endochondral bone or intramembranous bone, respectively (Abzhanov et al., 2007; Galea et al., 2021; Hall, 2015). Bone material is common to both types of bone and is a natural composite material that is present in the majority of vertebrate animals. The basic building blocks of bone material are collagen and hydroxyapatite crystals which form mineralized collagen fibrils (Reznikov et al., 2014; Weiner & Wagner, 1998), that combine with additional fibrils to form arrays, which can be organized to create different motifs such as woven bone, parallel fibered bone and lamellar bone (Francillon-Vieillot et al., 1990; Hall, 2015). Bone as an organ supports the body, protects internal organs and anchors muscles for locomotion (Francillon-Vieillot et al., 1990; Hall, 2015), and includes cells in tetrapods and basal bony fishes (but not in advanced teleosts), as well as blood vessels, nerves, non-collagenous proteins, and water (Currey, 2002; Reznikov et al., 2014; Rho et al., 1998; Wan et al., 2021; Weiner & Wagner, 1998).

Dermal bone, as the name implies, develops within the dermis of the skin and precedes the appearance of endochondral bone in the fossil record (Carter et al., 1998; Donoghue & Sansom, 2002). Dermal bone, together with enamel/enameloid and dentine, formed the elaborate exoskeleton of extinct vertebrates. Although some of these elements have been lost during evolution, dermal bone persists in living vertebrates (Hirasawa & Kuratani, 2015; Patterson et al., 1979). In extant vertebrates dermal bone forms a variety of skeletal elements including but not limited to scales, cranial bones, scutes, and fin rays (Francillon-Vieillot et al., 1990; Huysseune & Sire, 1998; Moss, 1968; Sire et al., 2009; Sire & Huysseune, 2003). Dermal elements are generally formed by woven bone and/or parallel fibered bone, although primary and secondary osteons (or lamellated bone) forming dermal skeletal elements have been reported (Atkins et al., 2015; de Buffrénil et al., 2015; Davesne et al., 2020; Johanson et al., 2022; Vickaryous & Hall, 2006; Witzmann & Soler-Gijón, 2010).

The endoskeletal long bones in the limbs of tetrapods are formed exclusively by endochondral ossification. Long bones grow in length from cartilage growth plates and increase in diameter by bone formation of periosteal surfaces. Endosteal and periosteal surfaces are first formed by woven bone which is later replaced by fibrolamellar, lamellar and/or osteonal bone (van Der Meulen et al., 1993; Enlow, 1962, 1966; Estefa et al., 2021; Francillon-Vieillot et al., 1990; Haines, 1942; Kronenberg, 2003; Locke, 2004; Singh et al., 1974; Talts et al., 1998). The cortex of long bones is the major load-bearing element and is formed almost exclusively from lamellar bone which may be in the form of osteonal or interstitial bone. Bone material possesses viscoelastic mechanical properties that provide long bones with resistance to bending and deformation under load, enabling their structural and functional roles (Currey, 1999, 2002; Martens et al., 1980; Morgan et al., 2018).

The sturgeon family of fish (*Acipenseriformes*) comprise 27 extant species that possess a skeleton with both cartilaginous and bony elements (Brownstein & Near, 2025; Hilton et al., 2011). In the fossil record sturgeons can be traced back for more than 200 million years (Bemis et al., 1997; Hilton et al., 2021), and molecular approaches estimate their divergence from the teleost lineage to have occurred between 345-390 million years ago (Du et al., 2020; Peng et al., 2007). This is close to the split between ray-finned fish and the lobe-finned fish line that eventually gave rise to tetrapods (Benton et al., 2015; Du et al., 2020). They are a basal ray-finned fish, characterized by a slow evolutionary rate and morphological stasis (Du et al., 2020). A morphological character that is a shared trait between sturgeons is a long and stout element located in the pectoral fin, called the pectoral fin spine.

Fin spines are formed by dermal bone and have a varied morphology across multiple fossil and extant non-tetrapod gnathostome lineages (Höch et al., 2021; Jerve et al., 2016; Kaatz et al., 2010; Kubicek et al., 2025). Early occurrences of paired and unpaired spines are found in extinct fossil jawed-fish such achantodians (Jerve et al., 2017; Watson, 1937), or in the stem-group osteichthyan *Lophosteus superbus* (Jerve et al., 2016). In basal and modern lineages of ray-finned fish, dorsal, pectoral, pelvic, and anal fin spines are also present (Meunier & Gayet, 2020; Price et al., 2015; Thacker & Near, 2025). In the context of the microstructure of fin spines, the appearance of concentric layers forming lamellar bone has been described in modern teleost dorsal (Kalish-Achrai et al., 2017) and pectoral fin spines (Abd-Elhafeez et al., 2024).

The pectoral fin spine (PFS) of the sturgeon is a robust dermal skeletal element that extends laterally and is located on the leading edge of the pectoral fin (Dillman & Hilton, 2014; Hilton et al., 2011). The PFS helps to maintain locomotor balance and stability for vertical maneuvering during swimming (Findeis, 1997; Wilga & Lauder, 1999), as well as for defense against predation and for contact to stabilize the position on the substrate during feeding (Findeis, 1993). The PFS is formed by the fusion of one or two initial pairs of pectoral fin rays (or lepidotrichia), a surrounding sheet of dermal bone, and a varying number of rays that are ultimately incorporated into the structure (Dillman & Hilton, 2014; Findeis, 1993, 1997; Hilton et al., 2011). The PFS has been used to estimate age in sturgeons (Bruch et al., 2009; Mosca et al., 2025), to provide insights into their ecology and life history (Allen et al., 2009; Bakhshalizadeh et al., 2023b; Neary et al., 2024), as a model for fin healing and regeneration (Allen et al., 2018), and to characterize marine pollution (Bakhshalizadeh et al., 2023a). However, a detailed description of the structure, composition and mechanical properties of the sturgeon’s pectoral fin spine in relation to the function it performs is lacking.

This study provides a detailed description of the structure of the PFS of the Russian sturgeon (*Huso gueldenstaedtii)*, and the mechanical properties and composition of the material from which it is formed. Using a multi-modal approach of imaging techniques (micro-CT, light microscopy, electron microscopy, polarized microscopy), mechanical testing (3-point bending, micro indentation), and composition analysis (FTIR, TGA, BMD), we show that the PFS has a complex microarchitecture, anisotropy, a high material toughness, and mineral composition that enable it to bear loads and to fulfill its role in locomotion and defense against predation. Additionally, we show dermal osteons in the PFS, which are structural features vital for load-bearing of tetrapod limb bones. These findings expand our knowledge of dermal bone and its ability to form structurally complex and mechanically competent skeletal elements for load-bearing, the functional and material similarities of dermal bone and endochondral bone, and increases our understanding of pectoral fin spines that are present in ray finned fish.

## Materials and methods

### Fish

Pectoral fin spines (PFSs) were harvested from 5-10-year-old male and 8-yearl-old female Russian sturgeons (*Huso gueldenstaedtii*) acquired from a commercial farm (Kibbutz Dan, Israel). Under farming conditions, male and female Russian sturgeons reach sexual maturity at ages of 3-4 years-old and 7 years-old, respectively (Hurvitz et al., 2008). Therefore, these specimens are adults, albeit at an early stage of adulthood. Pectoral fins are removed when dressing the carcass and were collected immediately after slaughter, placed on ice, and transported to the Laboratory of Bone Biomechanics (Koret School of Veterinary Medicine, Rehovot, Israel). Upon arrival, the fins were wrapped with gauze moistened with saline solution and stored at −20° C. Pectoral fin spines were isolated from the pectoral fins as required by thawing the pectoral fin at −4°C for 24 hours and then placing the fin in saline solution at room temperature for at least 2 hours. The PFS was dissected and cleaned of soft tissue, measured with a caliper (± 0.1mm) and the dimensions of the spine were recorded. In a preliminary study it was determined that the middle third of the PFS was the location that provide a good overall representation of the structure and sufficient material to prepare all the samples used in this study. This is similar to vertebrate long bone research where the mid-diaphysis is traditionally used (Prondvai et al., 2014). The middle third was defined as the region of interest and was harvested from all specimens used in this study using water-cooled rotary diamond saw (Isomet® low speed saw, Buhler).

### Fourier transformed infrared spectra (FTIR)

Sections harvested from the middle third of 3 spines were placed under an infrared lamp until dry and analyzed immediately. Once dry the scales were ground into a powder using an agate mortar and pestle. The ground spines were mixed with 2–3 mg of spectro-grade pure KBr, and pressed into a 5-mm diameter pellet using a Pike hand press (Pike Technologies, Fitchburg, WI, USA). The infrared spectrum of the transparent pellet was obtained using a Nicolet iS5 spectrometer (Thermo Scientific) at 4 cm ¹ resolution in the range of 4000 to 400 cm ¹.

### Thermogravimetric analysis (TGA)

A single cube (2mm x2mm x2mm) was harvested from the middle third of three PFSs and each cube was crushed to powder using a pestle and mortar (Jizerská porcelánka s.r.o., Desná, Czech Republic) until a fine powder was obtained. The powder was left in acetone overnight after which it was transferred to a platinum pan and placed into an 0 thermo gravimetric analyzer (TA Instruments, Waltham, MA). The sample was heated at a rate of 20°C per minute to 800°C under a nitrogen gas flow of 100 mL/min, using the Hi-ResTM technique. The temperature at which each component was lost and the relative weights of the water, organic, and inorganic (mineral) components of the pectoral fin spine were calculated using the TA Universal Analysis software (version 4.5A TA Instruments, Waters LLC, New Castle, DE).

### Reflected Light Microscopy

Reflected light microscopy images were acquired from thin (∼200μm) transverse and longitudinal sections prepared from the middle third of 9 PFSs. The middle third of the spine was first embedded in epoxy-resin and hardener (EpoThin ™ 2, Buehler, Il, USA) and then left to harden at room temperature for at least 24 hours. Sections were then cut in both the transverse and longitudinal planes. The surface of each section was ground and polished in a precision finishing machine (TechPrep, Allied High Tech Products Inc., Rancho Dominguez, CA, USA). Silicon carbide sandpapers (Buehler Metaserv, Buehler, Germany), with 400-, 800-, 1200-, 2500-, and 4000-grit were used for grinding, and a polycrystalline diamond suspension (3µm and 1µm) on a velvet cloth (Buehler Metaserv) was used for polishing. Images were captured using a reflected light microscope and a high speed digital microscope (Olympus® BX 51 microscope Olympus, Japan) with a dedicated camera. Panoramic views were generated by digitally stitching individual images using commercial software (Image Composite Editor, Microsoft Research, Redmond, WA).

### Polarized microscopy

Polarized microscopy images were acquired from thin (∼500μm) transverse and longitudinal sections prepared from the middle third of 3 PFSs. The middle thirds of the PFSs were demineralized in 10% EDTA for 4 weeks, after which they were embedded in paraffin and cut using a microtome (Leica RM2255, Germany). The resulting thin slices (∼2μm) were dehydrated in increasing concentrations of ethanol (70%, 80%, 95% and 100%), followed by three baths of xylene to deparaffinize them. Images were acquired using an LC-PolScope (CRi) analyzer mounted on a microscope (Nikon Eclipse 80i, Tokyo, Japan), and images of the collagen fibril orientations were captured with an LC-PolScope mounted on a microscope (Nikon Eclipse 80i, Tokyo, Japan).

### Back-scatter scanning electron microscopy (SEM)

Polished surfaces of the PFSs previously prepared for light microscopy, and fracture surfaces prepared in the transverse and longitudinal planes, were mounted on an aluminum stub with conductive carbon tape. Uncoated samples were imaged using the backscatter detector at low vacuum and 15 Kv EHT, 1024px resolution, working distance of 5mm (Phenom XL Desktop SEM, Phenom World BV, Thermo Fisher Scientific, Netherlands), and high resolution. In addition, polished and fracture surfaces were coated with carbon (5nm) and imaged using the backscatter detector in a field emission SEM at low vacuum, 20 kV EHT, 1024px resolution, working distance of 6.6mm (Sigma, Zeiss Oberlocken Germany).

### Micro-Computed Tomography (micro-CT)

One pectoral fin spine was scanned using a micro-computed tomography (micro-CT) scanner (SkyScan® 1174, Kontich, Belgium) to get a general overview of the three-dimensional structure of the spine. Due to space limitations in the micro-CT scanner the proximal flared end of the specimen was removed. The PFS was scanned through 180° of rotation, with a voxel size of 12.2 μm, an exposure time of 7000 ms, and a rotation step of 0.4°. The X-ray voltage source was set at 50 kV and the source current at 800μm. A 0.25mm aluminum filter was used to reduce beam-hardening effects.

The middle third of a PFS was cut and scanned through 180° with a voxel size of 13μm, an exposure time of 1679 ms, and a rotation step of 0.2°. The X-ray voltage source was set at 95 kV, the current at 105 μA, and an aluminum filter of 0.25 mm was used to reduce beam hardening. The data was reconstructed using commercial software (NRecon® Skyscan software, v 1.6.10, Bruker, Kontich, Belgium). The porosity of the spine in this location and the characterization of the network of canals within the PFS was determined using Dragonfly software v. 2022.2 (Object Research Systems, Montreal, Canada). A volume of 650 slices (length of ∼ 8.35 mm) was selected from the original stack. The slides were then segmented using the Segment Wizard trainer and refined by manual segmentation. After 26-connected component analysis, objects smaller than 27 voxels (equivalent to a 3×3×3 voxel region) were considered to be artefacts of scanning and excluded from the results. We then generated a threshold mesh, and obtained the mean and range values of the diameters of the neurovascular network. Subsequently, the Bone Wizard analysis tool embedded in the Dragonfly software was used to obtain the porosity of the middle third of the PFS.

Beams (1mm x 1mm x 15mm) were cut from the peripheral tissue on the dorsal (n=7), cranial (n=7), and ventral (N=7) aspects of the middle third of 7 PFSs using a water-cooled rotary diamond saw (Isomet® low speed saw, Buhler). A micro-CT scan (Skyscan® 1272, Kontich, Belgium) was acquired from each beam prior to mechanical testing. Beams were scanned through180° with a voxel size of 9 μm, an exposure time 613 ms, and a rotation step of 0.4°. The X-ray voltage source was set at 80 kV and the source current at 100 μA, and an aluminum filter of 0.25 mm was used. Once scanning was completed, phantoms of a known mineral density (0.25 and 0.75 g/cm3) were scanned using the identical protocol in order to determine the bone mineral density (BMD). All scans were reconstructed (NRecon® Skyscan software, v 1.6.10, Bruker, Kontich, Belgium) and analyzed (SkyScan’s CTAn v.1.15.9.0 software, Bruker, Kontich, Belgium). Porosity of the beam groups was obtained using the Bone Wizard analysis tool (Dragonfly software v. 2022.2) of approximately 10mm in length of each beam.

### Micromechanical testing

The twenty-one beams which were previously scanned were tested in 3-point bending in a custom-built micromechanical testing machine. The cranial aspect of each beam was loaded with the caudal aspect of the beam resting on two stationary supports 10 mm apart. The beam was immersed in saline during testing with the stationary anvil attached to the wall of the chamber and the moving anvil attached to a high precision linear motor (PI instruments, Germany) and a 110 N load cell (Sensotec, Columbus, OH). The force-displacement data generated when loading the beams was collected using custom written software (LabView, National Instruments, and Austin, TX) at 50 Hz. Stress and strain were calculated from the force-displacement graph using the following formulas:

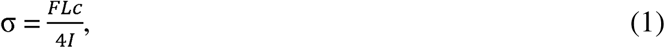

where σ is the stress (N/mm2), F is the force applied (N), L is the distance between the stationary supports (mm), c is half the thickness where the force was applied (mm) and I is the moment of inertia (mm4), and:

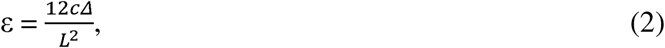

where ε is strain (%) and L is the vertical displacement of the beam at the point of load application (mm).

The following mechanical properties were calculated from the stress-strain graph: Young’s modulus of the material (E, GPa), yield stress, yield strain, max stress, max strain, ultimate stress, ultimate strain and work to fracture. The formulas to calculate I (moment of inertia) and E (Young’s modulus) were obtained from beam theory used previously on fish bone (Cohen et al., 2012).

### Microindentation

Cubes (3mm x 3mm x 3mm) were cut from the middle third of four pectoral fin spines using a water-cooled rotary diamond saw. For each cube the transverse and longitudinal surfaces were prepared for indentation by grinding and polishing the surface as described in for the preparation of samples for light microscopy. Samples were attached to a stub with wax and on each surface in each cube peripheral layered material and material concentrically orientated around voids were identified and indented (Micro-hardness tester, FM300, Future Tech Corp, Kanagawa, Japan). Indents were made with a Vickers micro-indenter tip (FM 100, Future-Tech©, NY, USA), using 5 gram-force (gF) loading force and 10 seconds dwell time. Subsequently, the Vickers hardness value (HV) was calculated using the ARS10k software. An indent was accepted only if the lengths of the two diagonals differed by less than 10%. Transverse and longitudinal surfaces were tested under both wet and dry conditions. Dry conditions were achieved by leaving the sample to dry at room temperature overnight, and wet conditions were ensured by submerging the cubes in saline solution for at least 12 hours prior to testing. Wet conditions were maintained during testing by periodically placing a drop of saline solution on the sample using a pipette.

### Statistics

Independent sample (pooled) t-tests were used to analyze the results of microindentation and micromechanical testing. Kruskal-Wallis test was used to analyze the results of beam porosity, the Steel-Dwass method for nonparametric comparison of all to significance beam between groups. In all tests p < 0.05 was considered statistically significant. JMP analysis software was used to perform the statistical analysis (JMP®, Version 19. SAS Institute Inc., Cary, NC, 1989–2025).

## Results

### Description of the pectoral fin spine (PFS)

The pectoral fin spine (PFS) and the anatomic planes defined to describe the structure are shown in Figure 1. The pectoral fin spine (PFS) of the adult Russian sturgeon (*Huso gueldenstaedtii*) is an elongated bone (see part 2: Composition of the PFS) which is located on the cranial/anterior aspect of the pectoral fin. The PFS is flared proximally where it attaches to the pectoral girdle, and becomes progressively narrower ending distally as a blunt point. A distally located, unfused section of the PFS, may persist in the mature fish.

**Figure 1.**
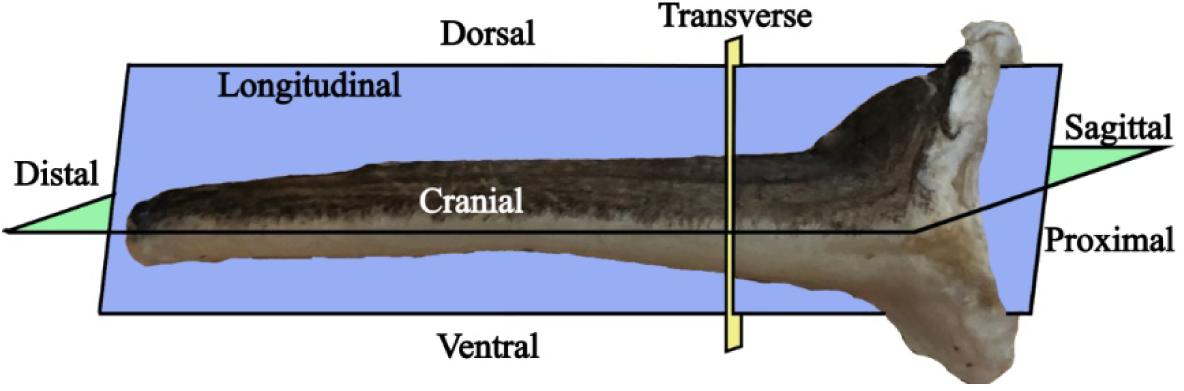
Photo of the right pectoral fin spine (PFS) showing its cranial aspect and the anatomical planes used to describe the structure. The transverse plane (yellow) divides the PFS into proximal and distal portions, the longitudinal plane (purple) divides the PFS into cranial and caudal parts, and the sagittal plane (green) divides the PFS into dorsal and ventral aspects.

### Composition of the PFS

#### Fourier transformed infrared spectrum (FTIR)

Fourier transform infrared (FTIR) spectra of the PFS (Fig. 2) is characterized by strong absorption bands of carbonated hydroxyapatite: 1453 cm^−1^ (carbonate ion v3), 901 cm^−1^ (carbonate ion v2) and 564 cm^−1^ (phosphate ion). There is also evidence of collagen: 1644 cm^−1^ (amide I), 1550 cm^−1^ (amide II), 1408 cm^−1^ (proline) and 1218 cm^−1^ (amide III). Comparatively, the mineral component of goat bone has peaks of 1467 cm^−1^, 874 cm^−1^ and 562 cm^−1^, respectively, while fresh pig cortical bone has collagen of 1653 cm^−1^, 1541 cm^−1^ and 1239 cm^−1^ (https://centers.weizmann.ac.il/kimmel-arch/infrared-spectra-library). The results of the PFS also correspond to other dermal bone elements of the sturgeon (*Infrared Spectra Library | The Helen and Martin Kimmel Center*, 2021; Milgram et al., 2023). Therefore, the PFS is a material with a composition equivalent of bone, as it has a spectra characteristic of collagen and hydroxyapatite (Lowenstam & Weiner, 1989).

**Figure 2.**
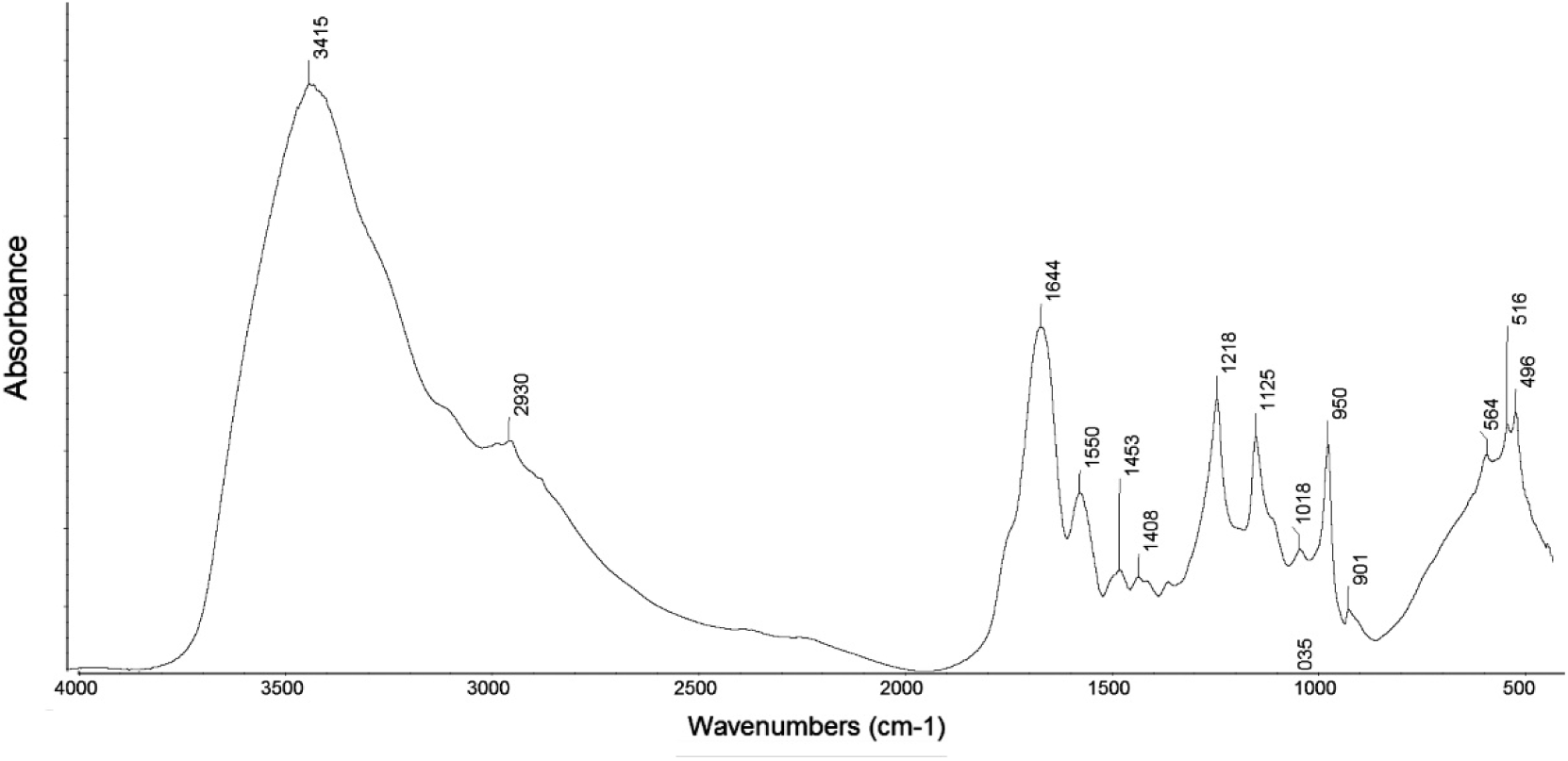
FTIR results of a fresh sample of the PFS. The peaks demonstrate the presence of collagen and hydroxyapatite, consistent with a bone-like composition.

#### Thermogravimetric analysis (TGA)

The PFS consists of an average (± sd) of 7.5% (± 0.9%) water, 31.8% (± 4.0%) organic material, and 60.7% (± 4.9%) inorganic material. These values are similar to those reported in literature for mammalian bone (Currey et al., 2001; Eastoe & Eastoe, 1954; Martin & Burr, 1989). Results show an initial decrease in weight at mean (± sd) temperature of 67.5°C (± 0.8°C) (n=3), which corresponds to the evaporation of the water bound to the bone matrix. The second decrease in weight occurs at a temperature of 333.4°C (± 1.9°C), which corresponds the “decomposition” phase, attributed to the combustion of the organic bone components, particularly collagen (Sup. Mat. 1).

#### Bone mineral density (BMD)

Bone mineral density (BMD) was calculated from the beams tested in 3-point-bending. The average BMD (± sd) of the PFS is 0.95g/cm^3^ (± 0.04g/cm^3^) of hydroxyapatite.

### Structure of the PFS

#### Micro-Computed Tomography (micro-CT)

The cranial, dorsal, and ventral aspects of the PFS form a convex surface (Fig. 3a) characterized by ridges orientated along the entire length of bone. Linearly arranged perforations (mean diameter ± sd: 20.20 µm ± 7.04 µm) that extend along the entire length of the spine are located in troughs between the ridges (Fig. 3b). The caudal aspect of the PFS is concave and forms a single longitudinally orientated groove along the length of the bone, with a similar linear arrangement of perforations of similar diameter (mean diameter ± sd: 22.46 µm ± 5.57 µm) (Fig. 3c, d). The mean proximal/distal length (± sd) of the PFS (n=25) used in this study was 9.0 cm (± 1.5 cm), however, in several specimens the last several millimeters of the distal end were separated from the PFS by soft tissue. In the middle third of the PFS the mean (± sd) cranial/caudal thickness is 0.9cm (± 0.2 cm), and the dorsal/ventral thickness is 6.5 mm (± 0.5 mm).

**Figure 3.**
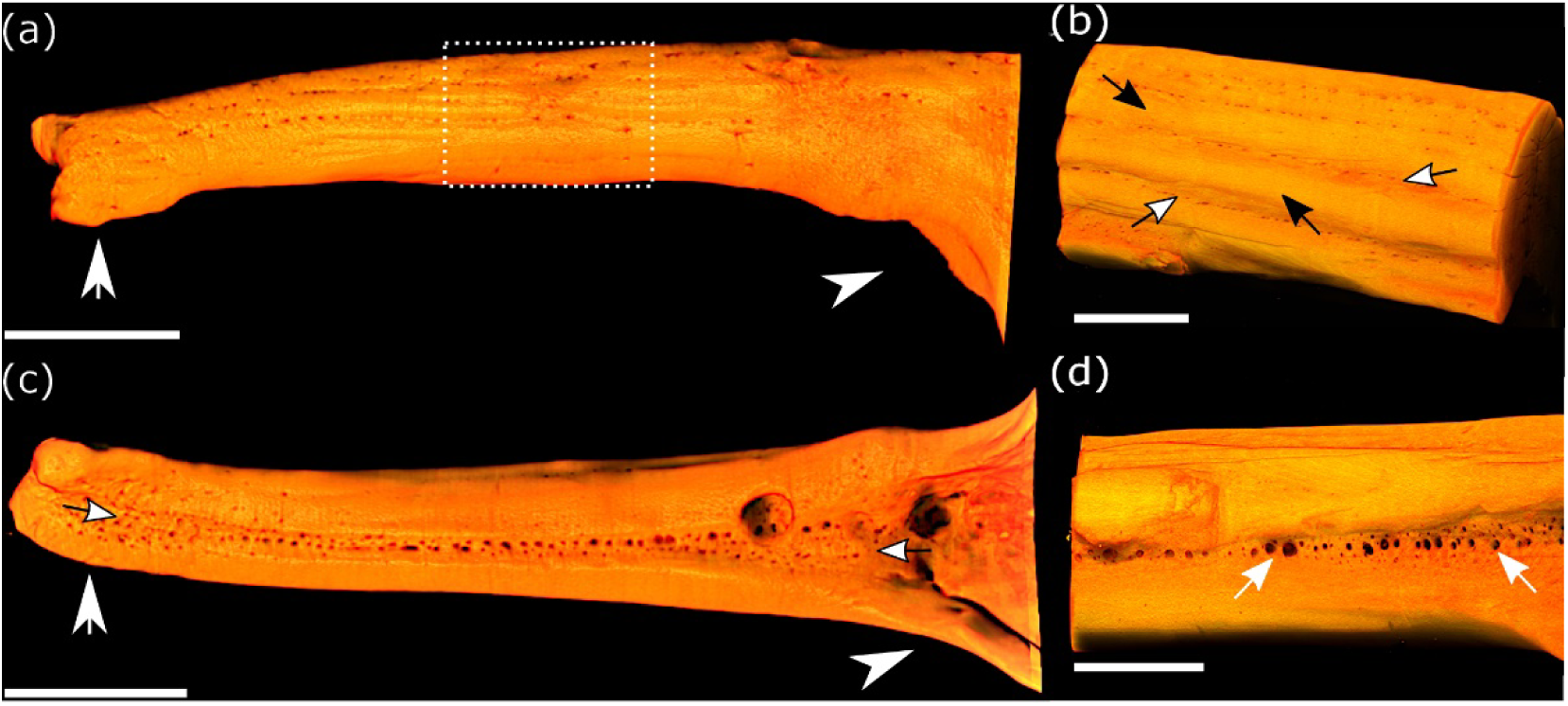
Micro-CT 3D reconstructions of the PFS. (a) Lateral aspect of a 3D reconstruction of an adult PFS generated from a micro-CT scan. The proximal end (white arrowhead) was cut prior to scanning, but the flare for attachment to the pectoral girdle is still visible. The irregular distal end (short white arrow) is due to an unfused point which was not included in the scan. The dotted rectangle in the middle of the PFS is shown at higher magnification in (b), where the longitudinally orientated ridges (black arrows) and troughs (white arrows with black stroke) which characterize the convex cranial surface of the PFS are seen. Perforations arranged linearly (white arrows with black stroke) along the entire length of the PFS are located in the troughs (white arrows with black stroke). (c) The caudal aspect of the PFS showing the proximal (white arrowhead) and distal ends (white short arrow) of the PFS with a single groove orientated along the entire length (white arrows with black stroke) of the spine. When the caudal aspect is viewed at a higher magnification (d) numerous linearly arranges perforations are seen in the depths of the trough along the entire length of the PFS (white arrows). Scale bars: a) 1 mm; b) 500 µm; c) 1 mm; d) 500 µm.

#### Reflected light microscopy

Two distinct regions can be seen on polished sections of the middle third of the PFS, prepared in the transverse plane, despite the lack of a clear boundary between them (Fig. 4a). A peripheral region is formed by circumferentially orientated layers forming an area of bone interrupted by radially oriented lineations (Fig. 4b). The central region is characterized by the presence of irregular round to oval voids of different diameters which are likely neurovascular cavities orientated at an angle to the plane of the image (Fig. 4c).

**Figure 4.**
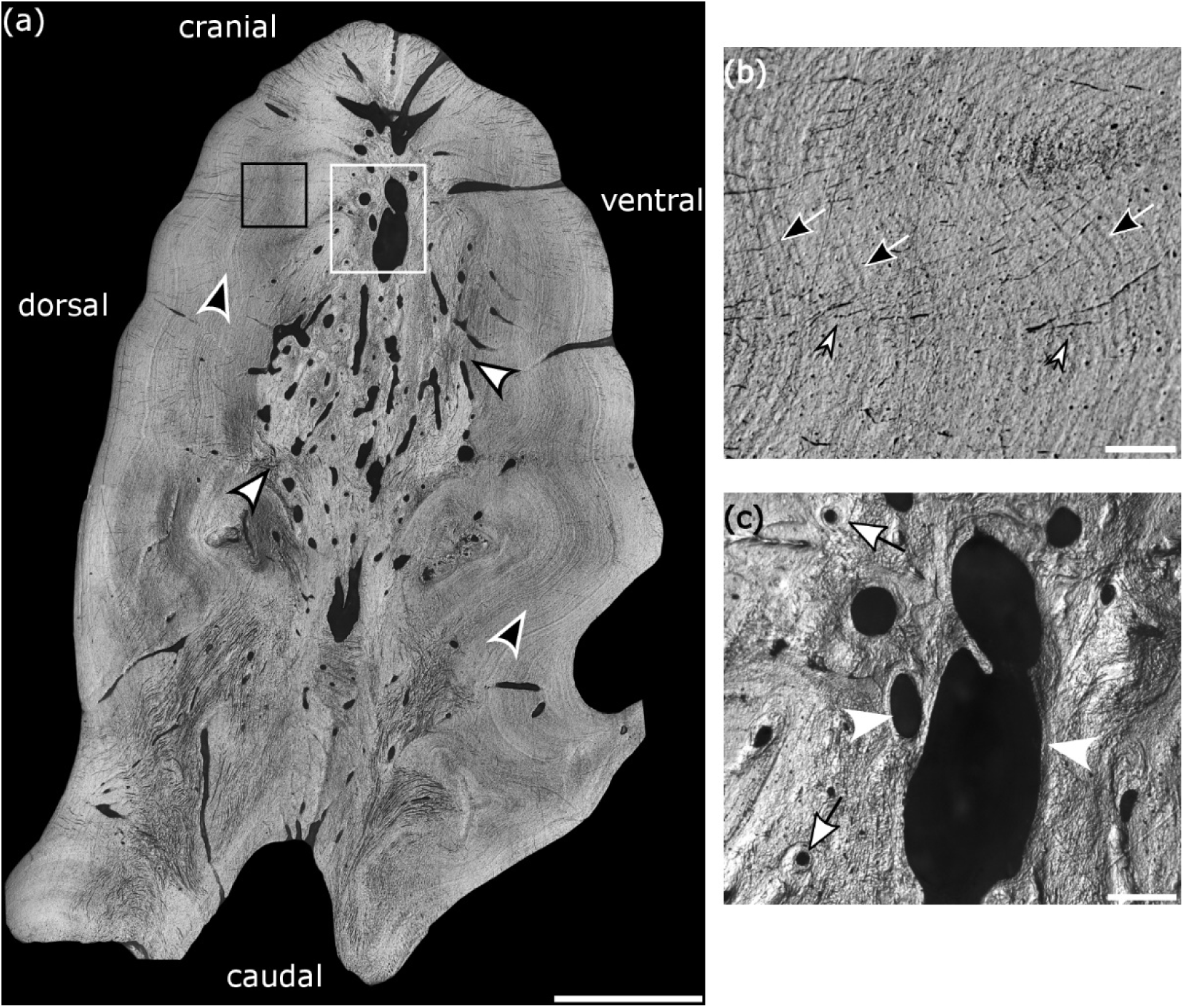
Transverse polished surfaces of the PFS using light microscopy a) Stitched reflected light microscopy image of the PFS prepared in the transverse plane showing a peripheral region formed by layers of bone material (black arrowheads with white stroke) surrounding a central region with irregularly round to oval voids of different diameters (white arrowheads with black stroke). A clear demarcation between the two regions is lacking. The black square in the peripheral region and the white square in the central region are shown at higher magnification in (b) and (c), respectively. (b) Circumferentially oriented layers of bone (black arrows with white stroke) and radially orientated lineations (short white arrows with black stroke) are seen more clearly at higher magnification of the peripheral region. (c) The presence concentric layers of material surrounding the larger diameter irregular round to oval voids (white arrowheads) and smaller diameter more circular voids (white arrows with black stroke) can be seen at higher magnification of the central region. Voids with very small diameters randomly distributed in the tissue are described in Fig. 5. Scale bars: a) 1200µm; b) 200µm; c) 50µm.

Two regions without a clear demarcation are also seen in polished sections prepared in the longitudinal plane, where lineations form the layering of the peripheral region (Fig. 5a, b). In this orientation the voids are also irregularly round to oval and are surrounded by concentric layers of material (Fig. 5a, c). Also visible are small voids with long and short mean diameters (± sd) of 9.4 µm (± 2.8 µm) and 6.4 µm (± 1.5 µm) respectively, and a mean area (± sd) of 61.1 µm^2^ (± 26.4 µm^2^). These dimensions are consistent with those of osteocytic lacunae in other sturgeon mineralized structures (Milgram et al., 2023) and we refer to these small voids as osteocytic lacunae (Sup. Mat. 2).

**Figure 5.**
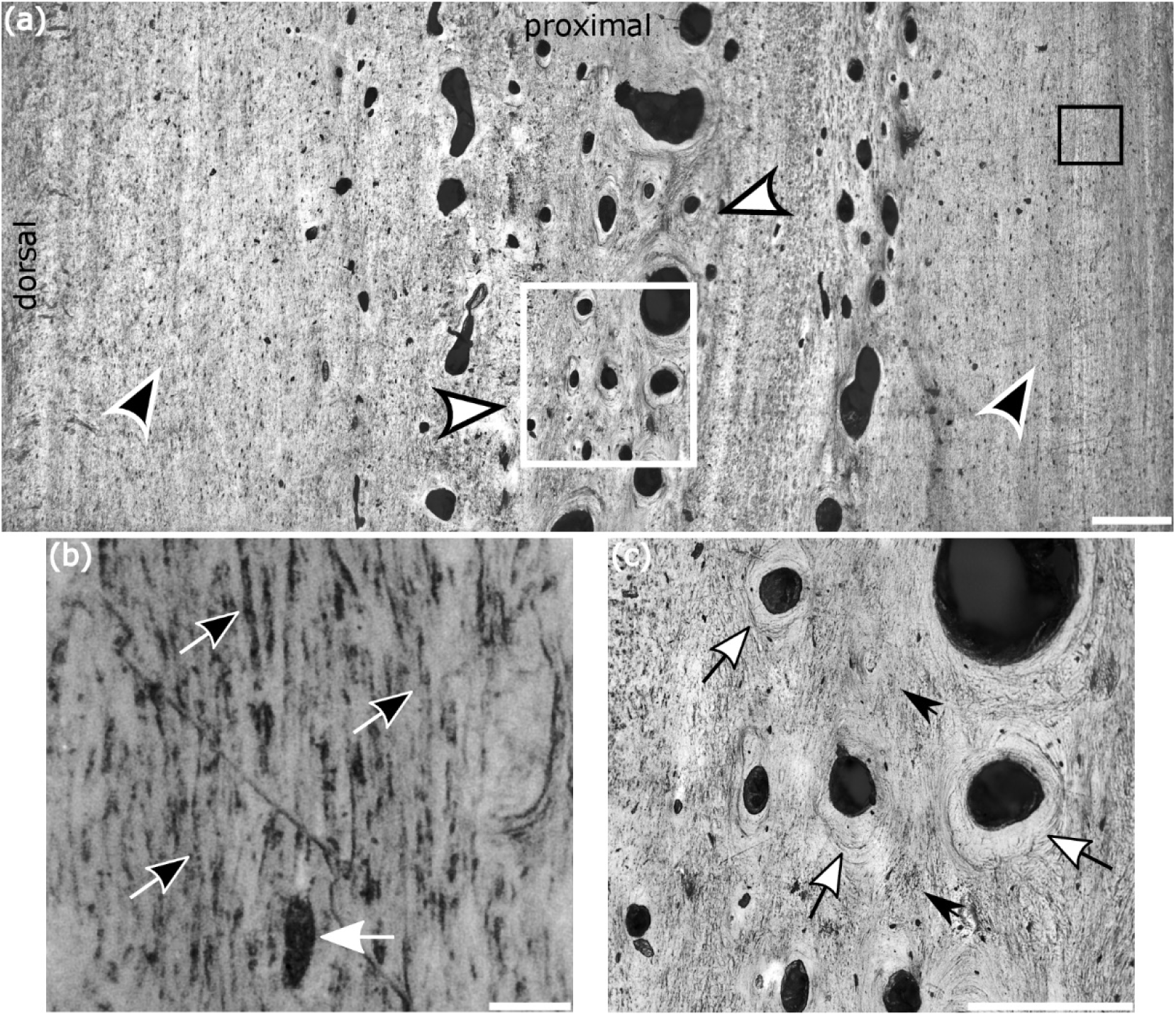
Longitudinal polished surfaces of the PFS using light microscopy. (a) Stitched reflected light microscopy image of the PFS prepared in the longitudinal plane showing a peripheral region formed by layers of bone material (black arrowheads with white stroke) surrounding a central region (white arrowheads with black stroke) with irregularly round to oval voids of different diameters. A clear demarcation between the two regions is not seen in this plane. The black square in the peripheral region and the white square in the central region are shown at higher magnification in (b) and (c), respectively. (b) At higher magnification the layering of the bone in the longitudinal plane is seen as vertical lineations and a void with the dimensions of an osteocytic lacuna (white arrow) is seen between two layers of material. (c) At higher magnification circular and oval voids are surrounded by circumferentially orientated layers of bone material (white arrows with black stroke), and are separated from each other by areas with no obvious structure (black arrows). Scale bars: a) 600µm; b) 15µm; c) 300 µm.

#### Back-scatter scanning electron microscopy (SEM)

Back-scattered electron detector images acquired from transverse and longitudinal polished surfaces of the peripheral region show that the vast majority of collagen fibril bundles (CFBs) are oriented in the same direction, with presumed CFBs oriented orthogonal to them (Fig. 6a, c). Fracture surfaces created in the transverse and longitudinal plane show most of the CFBs orientated in the plane of the image with a few presumed CFBs at an angle to the plane of the image (Fig. 6b, d). In all images CFBs are predominantly along the long axis (longitudinal plane) of the PFS. Presumed CFBs likely correspond to the radially orientated lineations seen in light microscopy. These CFBs have a mean thickness (±sd) of 2.7 µm (± 1.9 µm) and a mean length (±sd) of = 107.6 µm ± 100.7 µm) and are orientated normal to the external contour of the PFS.

**Figure 6.**
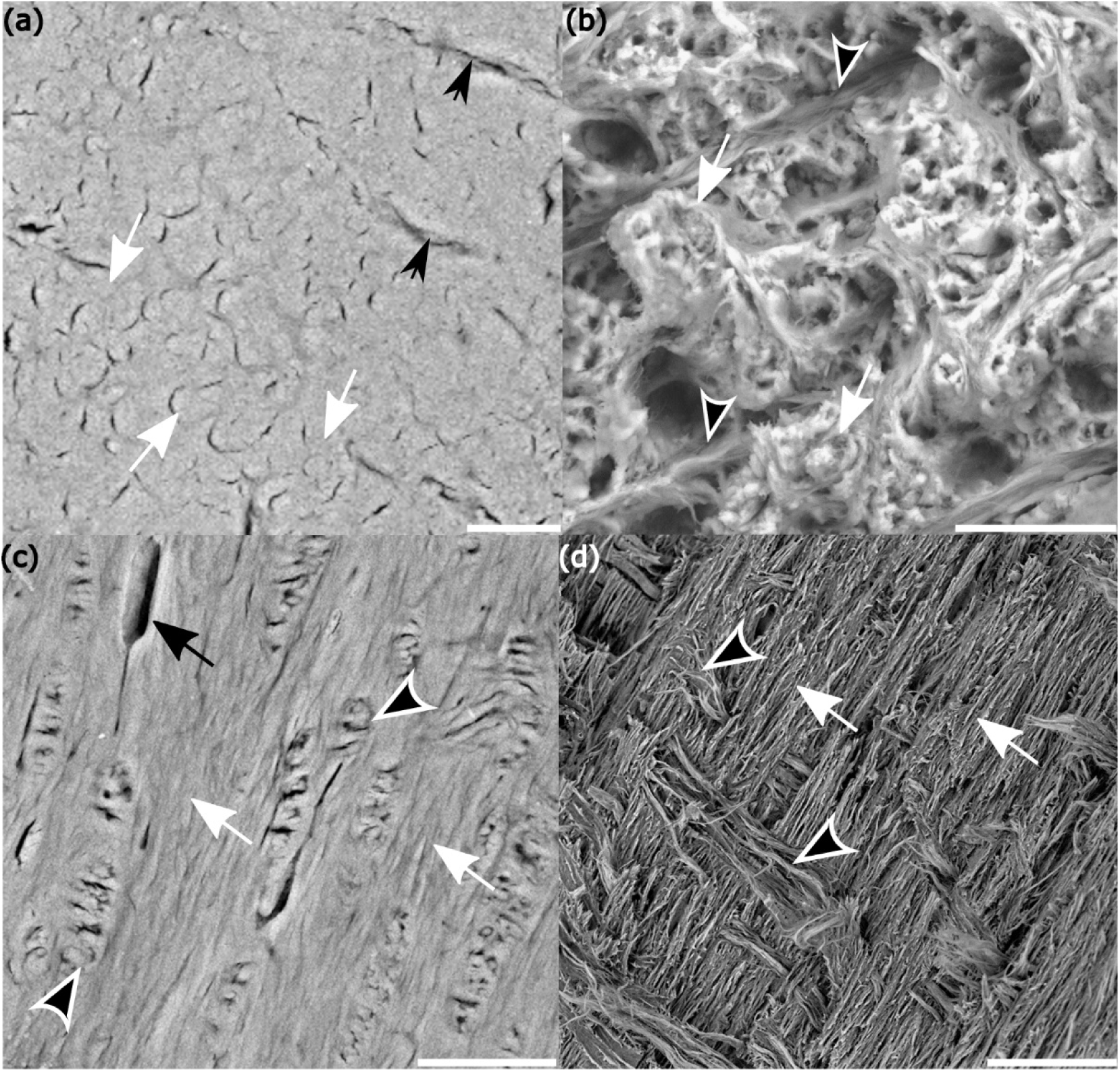
Backscatter electron detector images acquired from polished (a, c) and fracture (b, d) surfaces of the peripheral region prepared in the transverse and longitudinal plane. (a) Polished section prepared in the transverse plane with the collagen fibril bundles (CFBs) seen end on as they are orientated normal to the plane of the image and appear circular (white arrows) in this view. Radially oriented fine lineations (black arrows) in the plane of the image are also present. (b) Fractured surface prepared in the transverse plane with most of the CFBs orientated normal to the plane of the image (white arrows), however, presumed CFBs are also orientated in the plane of the image (black arrowheads with white stroke). (c) Polished surface prepared in the longitudinal plane with the majority of CFBs orientated in the plane of the image (white arrows) with assumed CFBs orientated at an angle to the plane of the image (black arrowhead with white stroke). An osteocytic lacunae with a canaliculus extending from it (black arrow) is also present. (d) Longitudinal fracture surface shows CFBs predominantly in the plane of the image (white arrows) with assumed CFBs orientated at an angle to the plane of the image (black arrowheads with white stroke). Scale bars: a) 20µm; b) 10µm; c) 40µm; d) 50µm.

Back-scattered electron detector images of polished and fracture surfaces of the central region show voids with concentric layers of bone surrounding them (Fig. 7a). The similarity of these structures to osteons described in vertebrate long bones is striking (Havers, 1691). This observation is strengthened by the presence of osteocytic lacunae between adjacent layers of bone material (Fig. 7a). Another feature in this region are filled voids surrounded by concentric layers of bone (Fig. 7b). These structures are reminiscent of sealed osteons present in human tibia and femora, as well as deer and horse limb bone (Congiu & Pazzaglia, 2011; Skedros et al., 2018). Hereafter, we refer to these structures as dermal osteons. Dermal osteons have a mean diameter (± sd) 187.3 µm (± 102.7 µm), while that of the central cavity (± sd) is 99.1 µm (± 71.5 µm). At high magnification, the material forming the concentric rings of the dermal osteons appears to form layers of CFBs (Fig. 7c). A fracture surface validates this observation as CFBs are similarly orientated both in and at an angle to the plane of the image (Fig. 7d). Additionally, the bone material between dermal osteons appears to be less organized than the layered bone of the dermal osteons (Fig. 7a, b).

**Figure 7.**
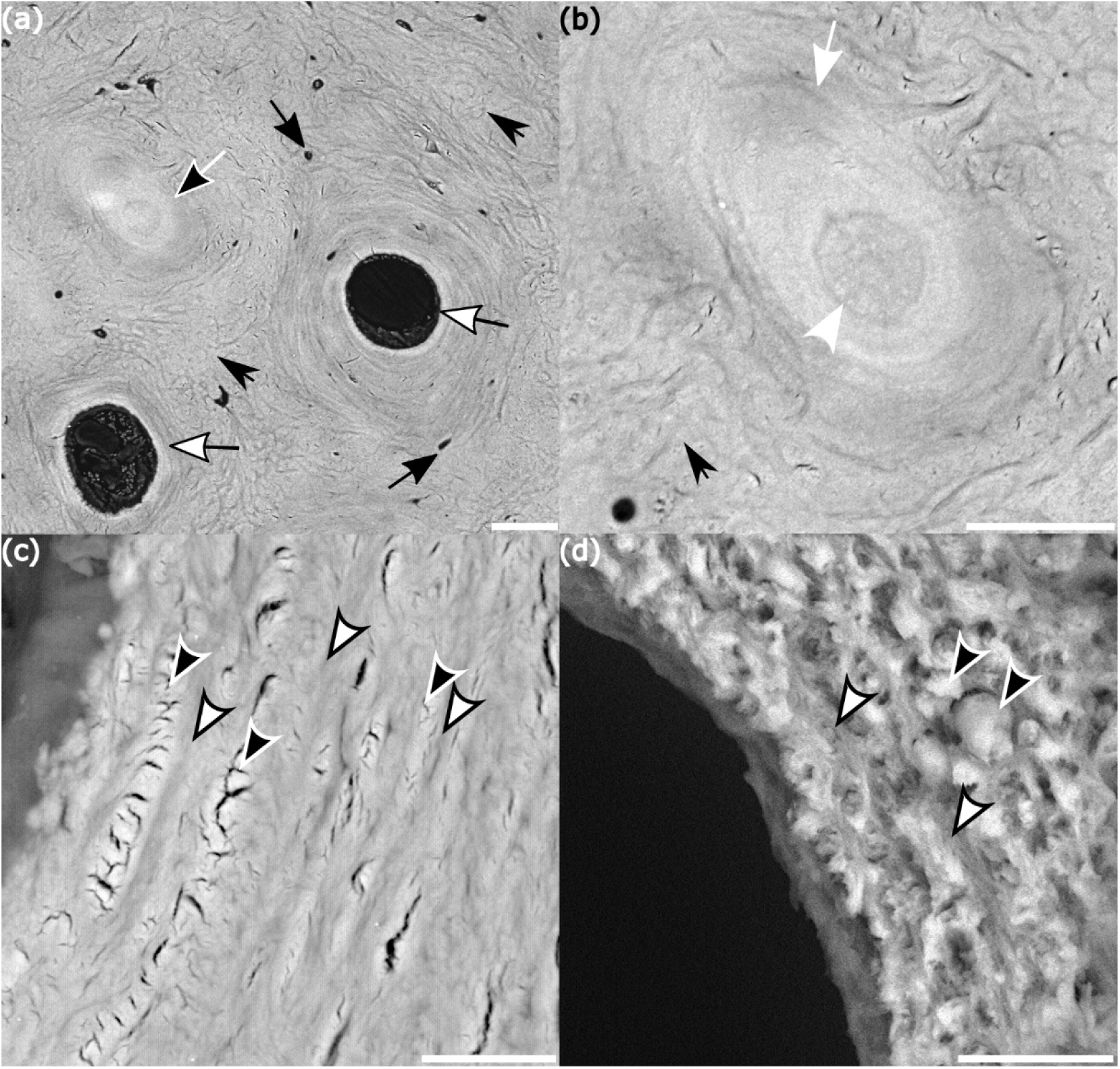
Backscatter electron detector images acquired from polished (a, b, and c) and fracture (d) surfaces of the central region prepared in the transverse plane. (a) Dermal osteons (white arrows with black stroke) are circular voids surrounded by circumferential layers of bone with osteocytic lacunae (black arrows) located between the circumferential layers of bone. Note that layering is not seen in the material between the dermal osteons (short black arrows) and we refer to this tissue as less organized tissue. The sealed dermal osteon (black arrow with white stroke) also seen in this image is reminiscent of the same structure in mammals, and is shown at higher magnification in (b). (b) A sealed dermal osteon is formed by layers of bone material (white arrow) forming concentric rings around an obliterated central canal (white arrowhead). Also visible is the less organized material (short black arrow) located between the osteons and sealed osteons. (c) At higher magnification the concentric layers of dermal osteons, appear to have CFBs orientated in two different orientations (black arrowhead with white stroke and white arrowheads with black stroke). This finding is confirmed on the transverse fracture surface of the osteon (d) where CFBs in the concentric layers are orientated in at least two different directions (black arrowhead with white border and white arrowheads with black border). Scale bars: a) 40µm; b) 30µm; c) 10µm; d)10µm

#### Polarized light microscopy

Polarized microscopy images of thin transverse and longitudinal sections confirms the predominant orientation of CFBs in the peripheral and central regions seen with the scanning electron microscope. CFBs in the peripheral region are orientated in a single predominant orientation, with no evidence of layering (Fig. 8a). This differs from concentric bone material forming the dermal osteons, which shows organized CFBs with different orientations surrounding a central canal (Fig 8b, c). In the central region (Fig 8d) the material between the dermal osteons is formed by CFBs orientated in different directions in the plane of the image. The material in this location corresponds to the less organized material described previously.

**Figure 8.**
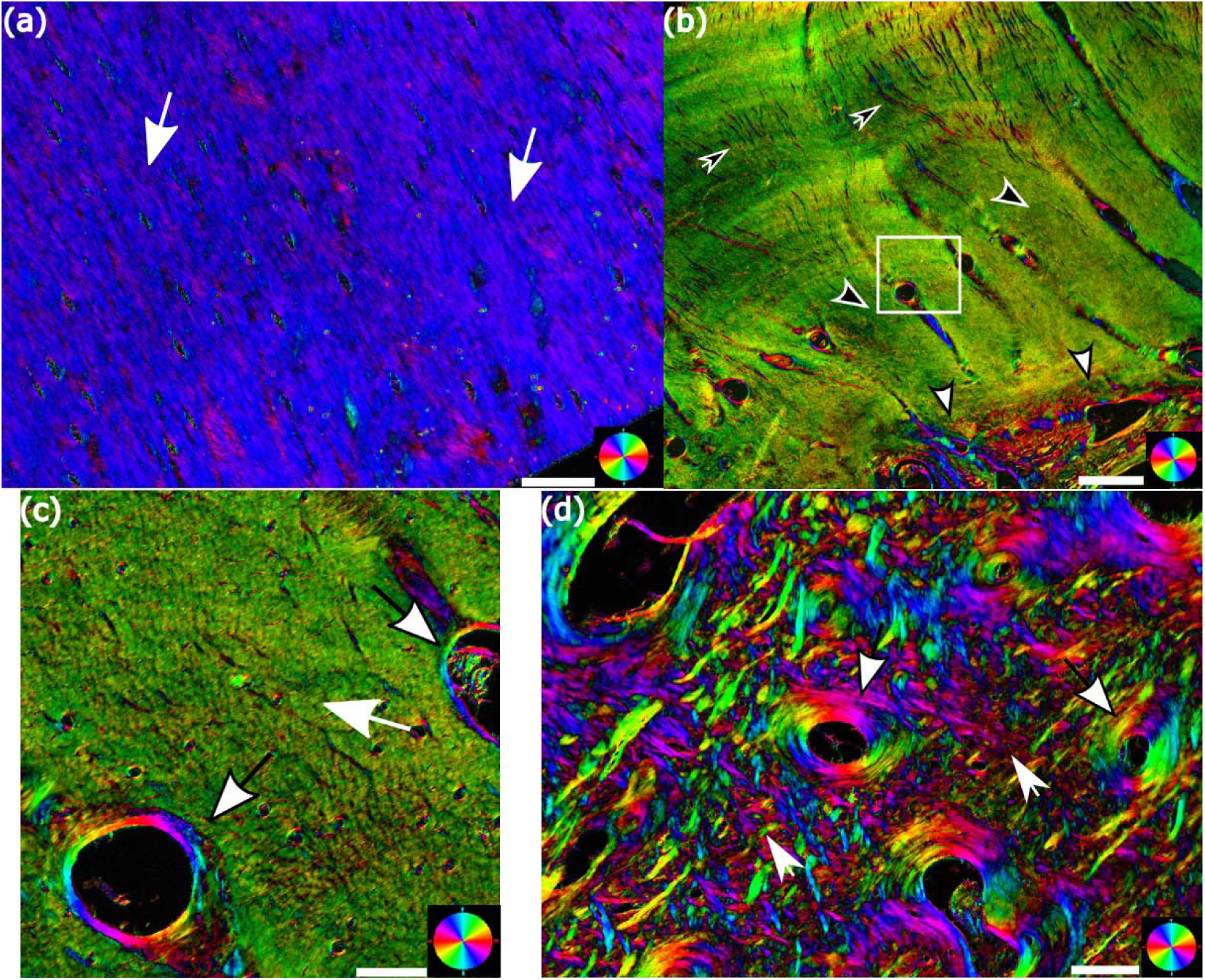
Polarized light microscopy images of the peripheral (a, b, and c) and central (d) regions of the PFS. (a) When viewing the peripheral region in the longitudinal plane the color (blue) is almost homogenous indicating that the CFBs (white arrows) are orientated predominantly in a one direction indicated by the color wheel in the bottom right corner. When viewing this region in the transverse plane (b) where the CFBs are orientated normal to the light source the tissue gets a green color (black arrowheads with white border) which is the color of the incident light passing through the section unchanged. Fine lineations are also seen (black arrows with white border). The transition between peripheral and the central regions (white arrowheads with black stroke) is seen in this section and white square surrounding a dermal osteon is shown at higher magnification in (c), where dermal osteons formed by CFBs orientated in different directions (white arrowhead with black border) surround the central cavities seen within the predominantly unidirectional CFBs (white arrow) which from the peripheral region. (d) In a transverse section of the central region, dermal osteons (white arrowheads), described in (c), are separated by material (short white arrows) with CFBs orientated in different directions in the plane of the image. This material corresponds to the less organized material described previously. Scale bars: a) 100µm; b) 300µm; c) 50µm; d) 50µm

#### Neurovascular canal network of the PFS

Segmentation of the voids at the center of the dermal osteons revealed an extensive network of neurovascular canals (average diameter = 140.08 µm) within the PFS. The largest diameter canal (Range: 362.60 µm - 466.33 µm) is located at the cranial aspect of the PFS and extends for the entire proximal-distal length of the PFS. This is the main canal of the PFS and gives rise to populations of smaller diameter canals which branch directly from it. Canals of intermediate diameter (Range: 259.08 µm – 362.60 µm) are orientated in the cranial-caudal direction and extend from the main canal to the caudal end (Figs. 9a, b). Small diameter canals (Range 52.03 µm – 259.08 µm) branch off the larger diameter canals and form a complex network of interconnections within the central region of the bone. In the peripheral region the small diameter canals, which branch directly off the main canal or are the terminal branches of the complex interconnections in the central region, are orientated radially towards the outer contour of the bone (Fig. 9a, b). It is likely that the perforations seen in the surface in the 3D reconstructions (Fig. 3) are the openings of these canals under the skin. This strengthens the claim that these canals are conduits for neurovascular bundles. The canals represent 2.82% of the porosity of the PFS.

**Figure 9.**
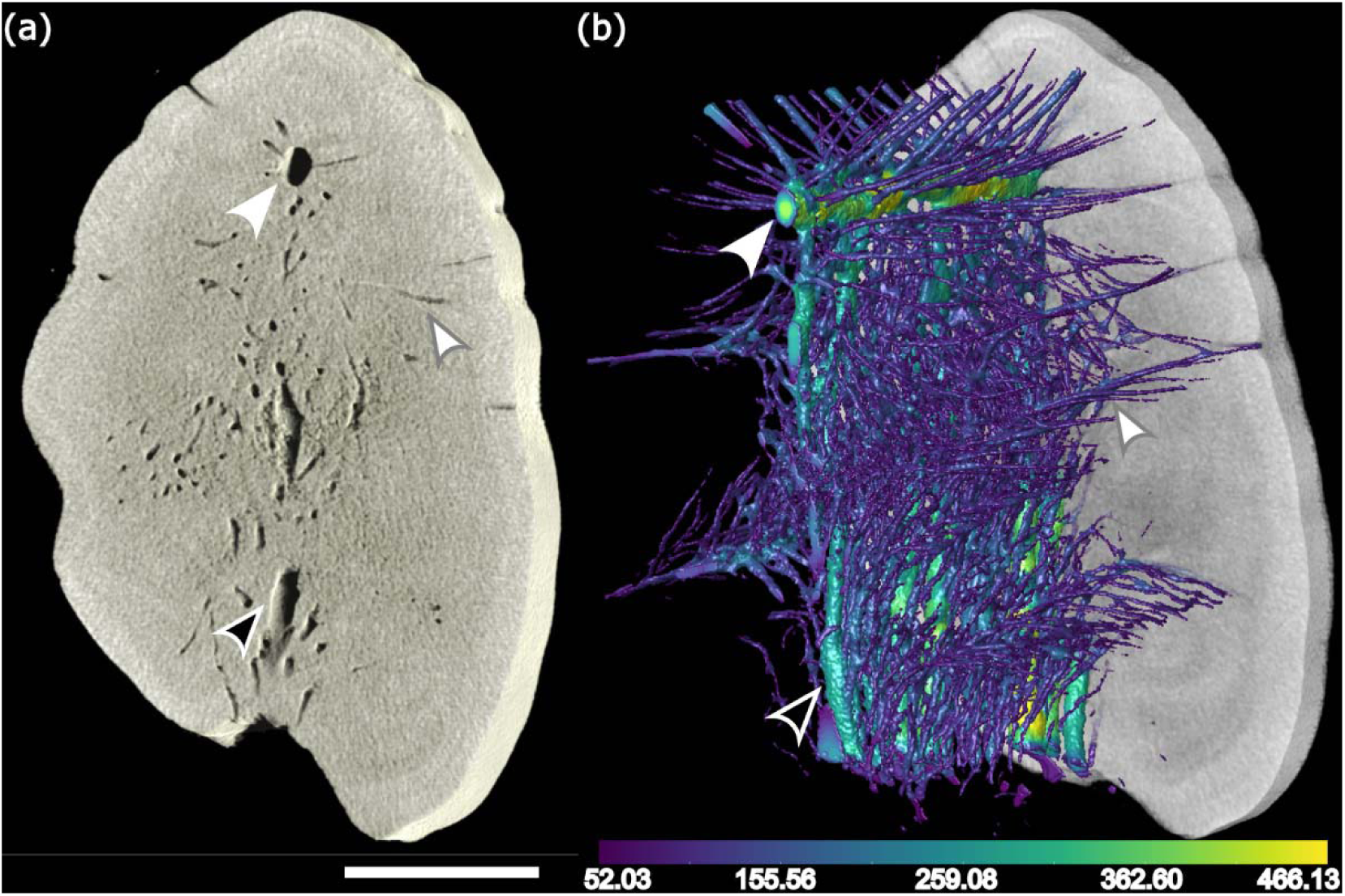
Transverse surface of a 3D reconstruction of a portion of the middle third of the PFS. (a) The main canal is orientated along the entire proximal-distal length of the PFS (white arrowhead). A secondary canal of a smaller diameter is orientated in the cranial-caudal direction (black arrowhead with white border). Smaller branches are in the radial direction are also seen (white arrowhead with grey border). (b) A 3D thickness mesh of the neurovascular network reveals the organization previously described in A). The color of the main canal and of the intermediate canals (yellow-green) indicate a larger diameter (white arrowhead and black arrowhead with white border). Smaller branches in multiple directions, including radially orientated canals (white arrowhead with grey border) have a different color code (dark green-purple). Scale: a) 1 mm

### Mechanical properties of the PFS

Twenty-one beams representing 7 different pectoral fin spines were tested in 3-point bending while submerged in saline solution and the results are shown in Table 1. An indication of the mechanical properties of the bone was determined from the stress/strain curves. The results show that the PFS has a mean (± sd) stiffness of 5.8 (± 1.5) GPa. The PFS also has a large post yield region, which translates into considerable deformation before fracture. The strain to fracture of the PFS (12.5 ± 4.9 %) and its considerable energy to fracture (14.8 ± 8.7 GPa) indicate that this bone has a remarkably high material toughness.

**Table 1.** Mechanical results of 3-point bending (mean ± standard deviation) of beams from the pectoral fin spine of the Russian Sturgeon

| Mechanical parameter | Result (mean ± sd) |
| --- | --- |
| *Young's modulus (E)* | $5.8 \pm 1.5$ GPa |
| *Yield stress* | $92.5 \pm 28.2$ MPa |
| *Yield strain* | $1.5 \pm 0.3$ % |
| *Maximal stress* | $137.3 \pm 37.9$ MPa |
| *Maximal strain* | $8.1 \pm 3.1$ % |
| *Fracture stress* | $92.0 \pm 34.2$ MPa |
| *Fracture strain* | $12.5 \pm 4.9$ % |
| *Energy to fracture* | $14.8 \pm 8.7$ GPa |

Micro-CT done prior to mechanical testing revealed that the while the dorsal and ventral beam were harvested from areas where the bone was uniform, the cranial beam included the main proximal-distal canal as well as many of the connected smaller diameter canals of the PFS. Porosity analysis confirmed that beams from the cranial aspect (16.69%) were more porous compared to beams from the dorsal (5.66%) and ventral (4.06%) aspects. Since no differences were seen in the mechanical properties when comparing the dorsal and ventral beams, we combined these groups and present our findings as a comparison between less porous and porous bone. (Fig. 10a).

**Figure 10.**
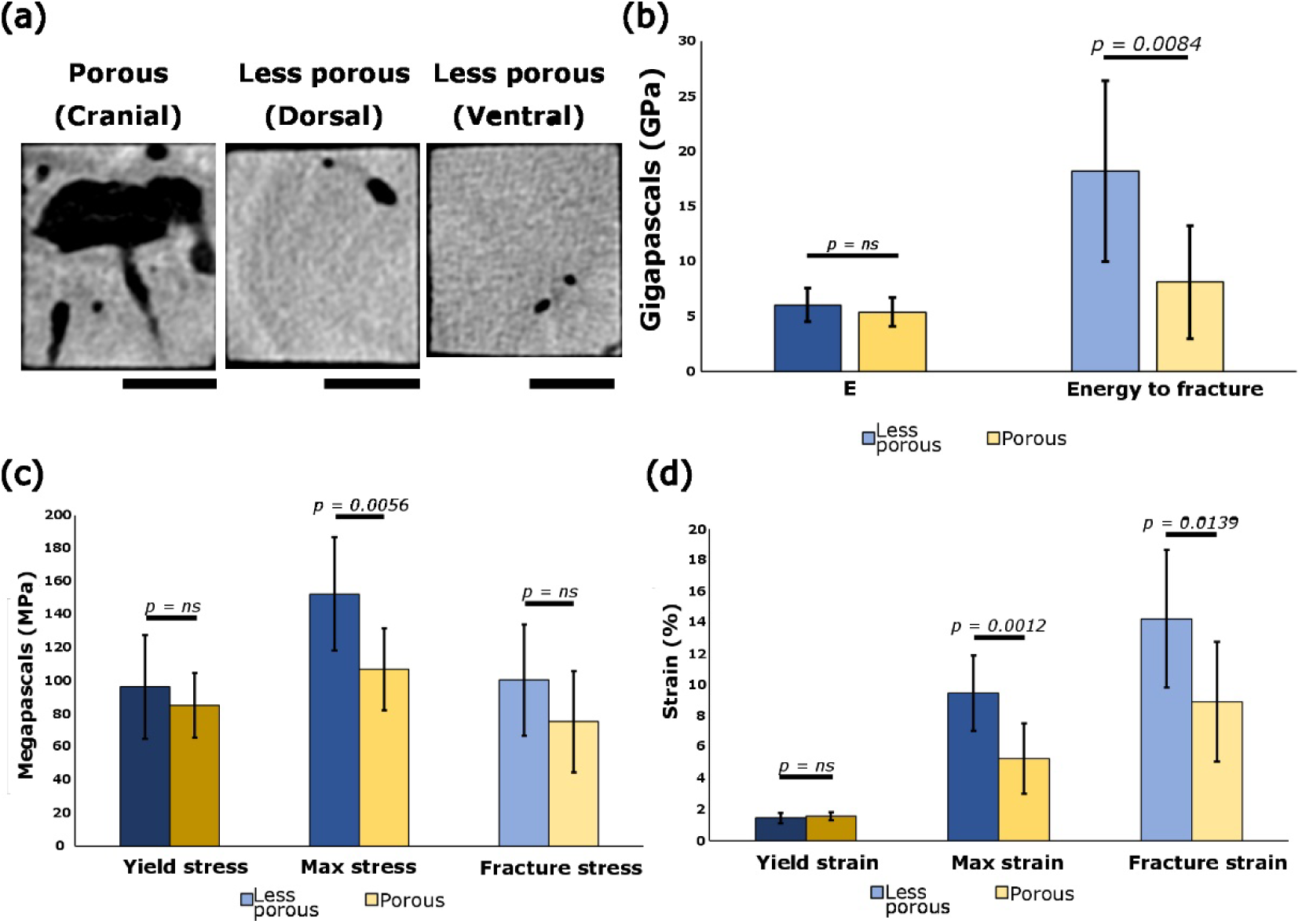
3-point bending results from less porous and porous beams of the PFS. (a) Image showing a cross section representative of the beams harvested from the cranial, dorsal and ventral aspects and tested in 3-point-bending. (b) Statistical analysis results comparing the less porous and porous bone of the PFS shows no significant difference in the Young’s modulus (E) but does show difference in energy to fracture. (c) Stress results are not different between the two groups of beams, except for maximum stress, and while no statistical difference was found in fracture stress, it was higher in less porous bone. (d) Significant differences were not seen in yield strain but were seen in the post-yield results (maximum strain and fracture strain) between less porous (peripheral) and porous (central) bone. Significant difference if p < 0.05

The Young’s modulus (E) was the same when comparing both groups (porous vs less porous), while the energy to fracture was significantly higher in the less porous bone material. This confirms that the material is the same in both locations, however, as there is more material dorsally and ventrally the energy to failure is higher (Fig. 10b). Pre-yield results were not significant between less porous bone and porous bone, but the decreased quantity of material due to the porosity affected the post-yield behavior of the beams. The results of the post-yield parameters of the less porous beams consistently exceeded those of the porous beam with stress at fracture the only parameter where significance was not reached (Fig. 10c, d).

Hardness value (HV) was determined from polished surfaces of cubes cut from the PFS. Bone from the peripheral region and the dermal osteonal bone were tested in the transverse and longitudinal planes under wet conditions and the results are shown in Figure 11. The peripheral bone material is anisotropic as it is significantly harder (p<0.0001) in the transverse plane when compared to the same material in the longitudinal plane. As the CFBs in this location are orientated predominantly in the proximal/distal direction, increased hardness would be expected when loading the CFBs parallel to their long axes. Similarly, the material forming the dermal osteons of the PFS is also anisotropic as the HV in the transverse plane is significantly (p=0.0015) harder than same material in the longitudinal plane. Interestingly, when comparing dermal osteonal bone to peripheral bone no significant difference was found in the transverse (p=0.5574) and longitudinal planes (p=0.5022). The results would suggest that under wet conditions, which are closer to physiological conditions, the two bone types have similar material properties.

**Figure 11.**
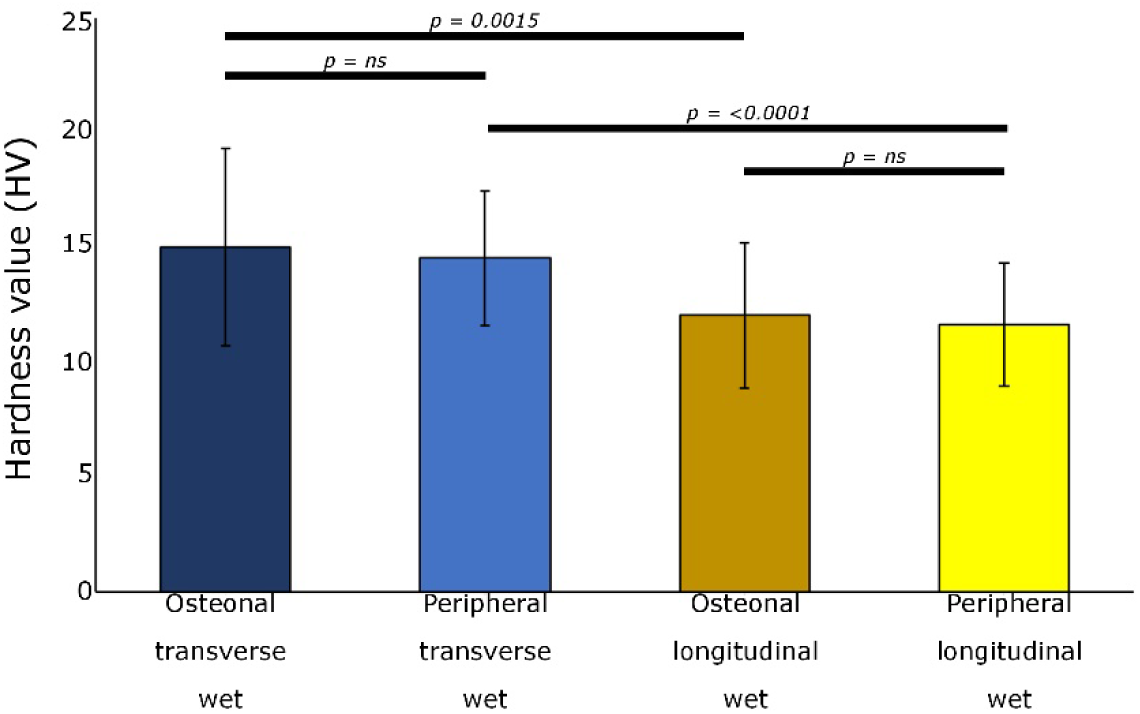
Hardness values (HV) of dermal osteonal bone and bone from the peripheral region tested in the transverse and longitudinal planes under wet conditions. Significant differences were seen when testing both types of bone in orthogonal planes. Osteonal and peripheral bone are both anisotropic as they were both significantly harder when loaded in the transverse plane, while no difference was seen between the two types of bone when loaded in the same direction.

The Young’s modulus (E) of the pectoral fin spine was calculated from the hardness data using the Oliver-Pharr formula (Oliver & Pharr, 1992). The E (mean ± sd) value was higher (p <.0001) in the transverse (proximal-distal) plane (5.89 GPa ± 1.1 GPa) compared to the longitudinal (cranial-caudal) plane (4.83 GPa ± 0.73 GPa).

## Discussion

In this study we describe structure, composition, and mechanical properties of the pectoral fin spine (PFS) of the Russian sturgeon (Huso gueldenstaedtii). At the micrometer scale the central region contains numerous dermal osteons with varying orientations formed by layered bone surrounding a central canal. The mineralized CFBs forming the material between the osteons is less organized and orientated in at least two orthogonal planes, however, large diameter CFBs were seen orientated along the long axis of the spine in the central region. In the peripheral region, at the micrometer scale, dermal osteons are fewer in number and radiate to the periphery in a more uniform manner. Although CFBs forming the tissue between the dermal osteons in this location are also orientated in orthogonal planes, the bulk of the material is orientated along the long axis of the bone in a single predominant direction. The orientation of the osteons suggests adaptation to multidirectional loading, whereas the alignment of the CFBs likely contributes to resistance against bending along the long axis of the spine.

The identification of two morphologically distinct regions within the PFS, characterized by differences in osteonal density and collagen fibril organization, suggests a functional partitioning of the spine at the organ scale. Such regional specialization is consistent with a structure adapted to complex loading conditions, where different structural elements contribute to mechanical performance. Notably, the finding that the vascular channels are surrounded by adjacent layers of layered bone and parallel-fibered bone, a configuration that likely supports both structural reinforcement and efficient load distribution around these channels, is similar to structural motifs described in fibrolamellar bone, a load-bearing tissue found in fast growing juvenile mammals, birds and vertebrates (Barrera et al., 2016). While the developmental and evolutionary relationships between these tissues remain unclear, this similarity raises the possibility that comparable mechanical demands have led to the emergence of analogous structural solutions in dermal and endochondral skeletal elements.

The magnitudes and directions of the loads experienced by the PFS are unknown. In mammalian long bones the osteons within the cortex are all aligned with the long axis of the bone and in the direction in which they are best able to resist the applied load (Cohen & Harris, 1958; Heřt et al., 1994; Tappen, 1977). The pectoral fin moves up and down as the sturgeon advances through the water (Wilga & Lauder, 1999) and based on the orientations of the osteons we speculate that the PFS is subjected to bending loads in three major directions (cranial-caudal, dorsal-ventral and proximal-distal).

The main canal of the PFS is located cranially on the midline of the spine and is orientated along the entire length of the long axis of the PFS. The main canal is surrounded by concentric layers of dermal bone forming a large dermal osteon, and as it is located close to the neutral axis of the spine, it is ideally positioned to resist the bending forces on the spine. The smaller diameter dermal osteons are orientated cranio-caudal and radially in different directions towards the external contour of the PFS. The radially orientated dermal osteons pass through the peripheral region, and penetrate the external surface of the spine where they are seen as pores on the external surface in the 3D micro-CT reconstructions. The distribution of dermal osteons likely indicates the direction of loading in addition to providing a rich blood and nervous supply to the overlying skin. In median and paired fin spines of acanthodians, networks of interconnected vascular canals have been described (Jerve et al., 2017). In mammalian long bone the canals of the Haversian system contain the neurovascular supply, are surrounded by osteonal bone, and are primarily orientated in the bone long axis, are connected by mostly transverse, shorter canals (Cohen & Harris, 1958; Currey & Shahar, 2013; Doube, 2022; Havers, 1691; Kim et al., 2015). It is important to note that the direction of Haversian systems can vary within and among bones, which might reflect the prevailing mechanical environment and loading mode (Cooper et al., 2004; Heřt et al., 1994). In addition,, the porous structure might contribute to its ability to absorb excessive energy (Quan et al., 2020). The lateral positioning of the PFS in the pectoral fin, and the fin participation in vertical maneuvering (Wilga & Lauder, 1999), suggest the loading of the PFS is complex which may explain the variety of orientations of the dermal osteons.

Dermal osteons are a major structural element of the central region of the PFS. In the PFS, the dermal osteonal diameter is well within the range reported in mammalian cortical bone although the central cavity is larger (Agerbæk et al., 1991; Cattaneo et al., 1999; Skedros et al., 2011; Zedda & Babosova, 2021). In vertebrate bone osteons can be either primary and secondary. Secondary osteons are the result of remodeling and generally overlap previously formed osteons. They are easily recognized by a cement line located at the periphery of the osteon (Enlow, 1962; Hall, 2015; Reznikov et al., 2014). Secondary osteons are rare in teleost and non-teleost fish bone, however they have been reported in anosteocytic billfish (Atkins et al., 2014), salmon (Witten & Hall, 2002), zebrafish (Weigele & Franz Odendaal, 2016) and horse-eye jack fish bone (Smith-Vaniz et al., 1995). We did not observe a structure similar to a cement line or any overlapping of dermal osteons in any of the specimens we examined. However, the oldest fish examined in this study was 10 years old and as the reported life span of sturgeons is up to 100 years (Hung, 2017), and we cannot exclude the possibility that remodeling occurs in the sturgeon at a later age. Interestingly, images captured at high magnification suggest that dermal osteonal bone is formed by CFBs with at least two distinct orientations, a feature in mammalian bone which contributes to crack deflection and enhanced toughness under mechanical loading.

Sealed osteons were first described in mammalian long bones in 1853 (Tomes & Morgan, 1853). Recent studies have suggested that they are part of the adaptation of bone to intracortical blood flow dynamics (Congiu & Pazzaglia, 2011) or are associated to the closing region of the cutting cone in secondary osteons (Skedros et al., 2018). Regardless of the physiological reason for their existence, they are not considered a pathological finding (Kornblum & Kelly, 1964; Skedros et al., 2018). The presence of sealed dermal osteons in the PFS (Fig. 12a-c) is noteworthy. Intracortical blood flow dynamics are beyond the scope of this study, and the absence of evidence of bone remodeling suggests that sealed osteons in the PFS are unlikely to be associated with cutting cone activity. Their occurrence in sturgeon bone, however, suggests that sealed osteons are a structural feature of load-bearing bone across many vertebrate lineages.

**Figure 12.**
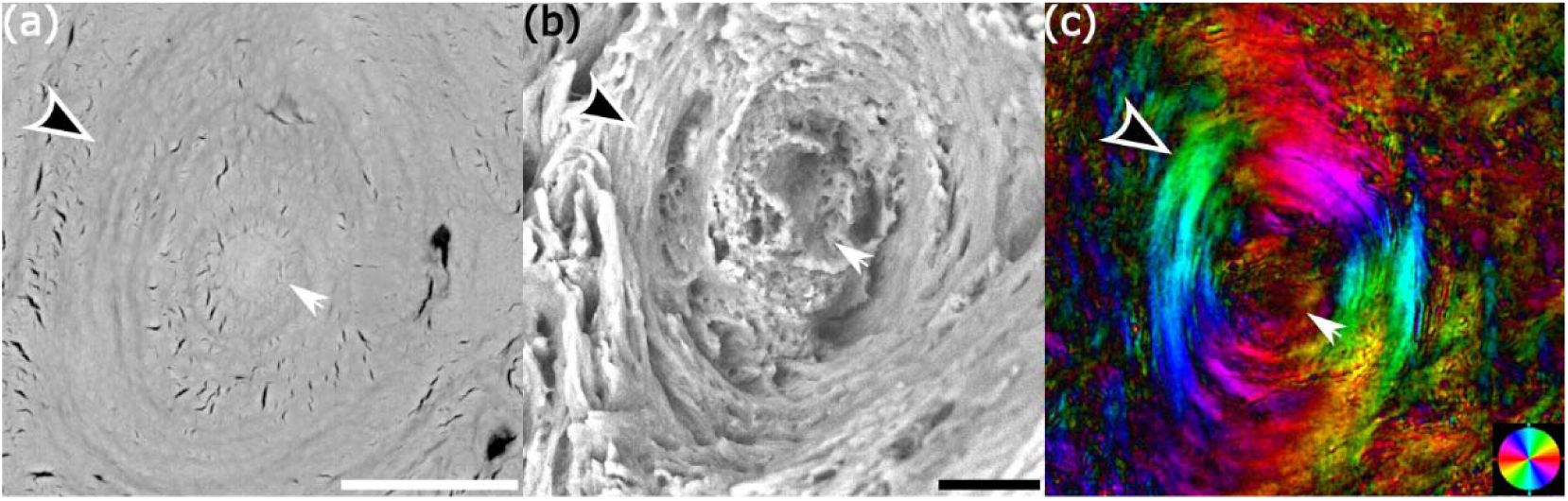
Backscatter electron detector images of a polished (a), and a fracture surface (b), of a sealed osteon prepared in the transverse plane, and polarized microscopy image (c), of the same structure in prepared in the same plane. Circumferentially arranged layers of bone (black arrowhead white stroke) seen surrounding a sealed central canal (white arrow) are shown in all images. Scale bars: a) 30µm; b) 15µm.

Peripheral bone of the PFS appears to be layered, but the bulk of the material is formed by parallel fibered bone. Mineralized collagen fibril bundles are predominantly orientated in one direction, parallel to the bone long axis of the spine where it likely increases the stiffness and resistance to bending along the principal loading direction. Parallel fibered bone is the most common type of bone found in teleost and non-teleost species (Davesne et al., 2018; Hughes et al., 1994; Qu et al., 2013; Raguin et al., 2020; Scheyer et al., 2014). The majority of bone material in developing teleost fish is parallel fibered bone, however, at the stage of development woven bone is also present (Weigele & Franz Odendaal, 2016). In mature teleost fish, parallel fibered bone remains the most abundant and may appear with layered bone (Apschner et al., 2011). In the PFS, there are presumed CFBs oriented at an angle to the long axis of the spine, and it remains to be seen if the material forming the peripheral bone has a layered motif at the nanoscale.

Mechanically, pectoral fin spines resist high bending forces and maintain toughness and strength allowing them to fulfill roles in defense and locomotion (Ferrón et al., 2021; Quan et al., 2020). In general, the PFS has a low Young’s modulus (E) and a remarkably high toughness, both of which are characteristic of fish bone (Atkins et al., 2014; Clifton et al., 2008; Cohen et al., 2012). A higher strain to fracture has been reported in anosteocytic billfish bone (bone (4% - 5%), compared to mammalian bone (2% - 3%) (Atkins et al., 2014), and in pectoral fin spines of other ray finned fish (Quan et al., 2020). Tissue-level hardness measurements demonstrate comparable hardness between peripheral parallel-fibered bone and dermal osteonal bone. The similarity in the mechanical properties of the bone tissue in the two regions is further supported by the similarity in elastic modulus (E) derived from the testing of the beams, indicating that both regions contribute comparably and effectively to load-bearing performance. In the spine, anisotropy was found to be a characteristic of dermal osteonal and peripheral bone. Anisotropy is a well-established mechanical property of mammalian long bones (Bonfield & Grynpas, 1977; Saha, 1973), evidence of mechanical function adapted to direction of loading (Nakano, 2015), and in the PFS it directly reflects the preferential orientation of CFBs and osteons observed.

The similar elastic moduli of the two regions of the PFS likely contributes to the absence of a sharply defined boundary between them. In biological structures composed of adjacent tissues with markedly different moduli, stress concentrations at the interface are often mitigated through the presence of intermediate material gradients or specialized interface architectures that facilitate load transfer and reduce fracture risk. In contrast, in the PFS, the adjacent regions exhibit similar deformation behavior within the elastic region of the stress–strain curve, which is consistent with strain compatibility and efficient load transfer without the need for a pronounced transitional layer, and may contribute to enhanced structural integrity under bending loads. While the architectural differences between these bone tissues would be expected to influence their mechanical properties, similarities in mineralization and/or water content may partially explain the observed the equal mechanical behavior (Cohen et al., 2012; Currey, 2002; Rho et al., 1998). Taken together, these structural and mechanical characteristics indicate that dermal bone in the PFS forms a mechanically competent, load-bearing element, exhibiting key functional and material similarities to endochondral long bone.

Bone mineral density of the PFS (0.75–1.0 g/cc) is within the range of amniote bone, it is similar the BMD values of dorsal fin spines in *O. aureus* and *C. carpio,* but lower when compared to mature mammalian bone – range: 1.20–1.30 g/cc or higher(Ashique et al., 2022; Atkins et al., 2014; Currey, 2002; Fletcher et al., 2018). It is important to mention, however, that specimens in our study were young adults that had probably not reached peak bone mass, and it would be expected for them to increase their BMD as they age. In contrast, ash content of the PFS is above 60%, even though cellular fish bone tends to have a low ash content (∼50%), which is related to the porosity of the lacuno-canalicular system (Cohen et al., 2012; Currey, 2008; Dean & Shahar, 2012; Horton & Summers, 2009; Toppe et al., 2007). In this regard the bone forming the PFS is similar to mammalian long bone (Clifton et al., 2008; Dean & Shahar, 2012). These compositional characteristics reinforce the interpretation that the PFS achieves a level of material organization and mineralization consistent with load-bearing skeletal elements, further supporting its functional equivalence, in key respects, to endochondral bone.

The gross morphology of the pectoral fin spine of the Russian sturgeon is similar to that of catfish (Vanscoy et al., 2015). Catfish, like sturgeons, are benthic feeder, and use the pectoral spines as a mechanism of protection. In extinct and extant actinopterygians and tetrapods, bone sculptures or ornamentations are present in the surface of dermal skeletal elements and include the formation of pits or holes, ridges and troughs (Clarac et al., 2017; Hill, 2006; Lundberg & Aguilera, 2003; Witzmann et al., 2010). Ornamentation is also characteristic of the gross morphology of catfish (order: Siluriforme) pectoral fin spines (Vanscoy et al., 2015). It is then not surprising that we also found these elements present in the external surface of the PFS. Especially intriguing is the relationship between the pits or holes located in the troughs of the external surface and the neurovascular canals. The pits or holes are thought to be closely associated with vascularization (Janis et al., 2012; Witzmann et al., 2010), and could have been used by early tetrapods to buffer acidosis and maintain it for longer periods while on land (Janis et al., 2012), or as a regulator of body temperature (Seidel, 1979). While we are not able to determine their true function(s), our observations support the hypothesis that these features are associated with vascularization.

The structural organization of the PFS, comprising parallel-fibered and layered bone associated with vascular channels, shares notable structural similarities and mechanical features such anisotropy, are consistent with load-bearing function with vertebrate fibrolamellar bone, despite the dermal origin of the PFS (Almany Magal et al., 2014; Barrera et al., 2016; Seto et al., 2008). While these similarities may reflect comparable mechanical demands leading to convergent structural solutions, the extent to which they represent conserved developmental or evolutionary features remains unresolved. Resolving this question will require investigation at the nanoscale, where differences in collagen-mineral organization may distinguish between true homology and functional analogy.

One of the main limitations of this study is the limited knowledge of genes or genetic pathways that may be shared between the sturgeon and other non-tetrapod gnathostomes where pectoral fin spines occur. We also are not aware of the cellular or structural features of the PFS during growth and development. This is a further limitation of this study; however, we are currently addressing this issue. A definitive method to classify dermal osteons as primary or secondary osteons is an additional limitation. Osteons are classified by morphological differences, however, documentation of fundamental differences in the material forming the different osteons is lacking. Structure of the circumferential layered bone forming secondary osteons in mammals has been described at the nanometer scale, however, bone forming primary osteons has not been characterized at this length scale. A better understanding of the differences between primary or secondary osteons in mammals at the nanometer scale would provide context for the classification of dermal osteons.

We have documented the structural, mechanical and compositional characteristics of the pectoral fin spine (PFS) of the Russian sturgeon. The evidence here presented (a complex microarchitecture formed by dermal osteonal bone and parallel fibered bone, high material toughness, considerable mineral ash content, anisotropy) show traits that allow the PFS to sustain high bending loads and fulfill its function for locomotion and defensive purposes. The presence of osteons in the sturgeon’s PFS, a key feature of tetrapod bone crucial for load-bearing, adds a noteworthy descriptive element. Additionally, we describe features (dermal ornamentation, a neurovascular network) that are also found in other pectoral fin spines of non-tetrapod jawed vertebrates. These findings expand our understanding of dermal bone and its ability to form structurally complex and mechanically competent load-bearing elements. They also provide evidence of functional and material similarities between dermal and endochondral bone and further inform our knowledge of pectoral fin spines in basal fish.

## Acknowledgments

The authors would like to acknowledge Prof. Steve Weiner for his advice and use of the Scanning Electron Microscope. We would also like to thank Prof. Ron Shahar for his assistance in the analysis of the mechanical testing results and polarized microscopy interpretation. We thank Prof. Rivka Elbaum for support with the Polarized Light Microscope. We also appreciate the help of Dr. Jeny Kertsnus-Banchik for support with the Thermogravimetric analysis experiment, and Dr. Tally Kossovsky for support with micro-CT. We thank Dr. Maïtena Dumont for her help with sample preparation and instruments, and fruitful discussion regarding the experiments and analysis. Many thanks to Dr. Orly Lewis, for her support. Special thanks to Dr. Avshalom Hurvitz at the Sturgeon Fishery, Kibbutz Dan, Israel, for generously providing access to the samples that were used in this study. This work was supported by funding from the Israeli Science Foundation (grant # 1751/22).

## Author contributions

Esteban Marroquín-Arroyave: Formal analysis, Data curation, Investigation, Validation, Visualization, Writing – Original draft, Writing- Review and editing. Josh Milgram: Conceptualization, Data curation, Funding acquisition, Investigation, Methodology, Project Administration, Resources, Supervision, Validation, Visualization, Writing- Review and editing.

## Data and materials availability

All data is available upon request directed to the corresponding author.

## Supplementary material

Please see additional document

## References

1. Abd-Elhafeez, H.H., Massoud, D., Mahmoud, M.S., Abdellah, N., Salah, A.S., Mohamed, N.-E., et al. (2024) Microstructural architecture of the bony scutes, spine, and rays of the bony fins in the common pleco (Hypostomus plecostomus). International Journal of Veterinary Science and Medicine, 12, 101–124. Available from: 10.1080/23144599.2024.2374201

2. Abzhanov, A., Rodda, S.J., McMahon, A.P. & Tabin, C.J. (2007) Regulation of skeletogenic differentiation in cranial dermal bone. Development, 134, 3133–3144. Available from: 10.1242/dev.002709

3. Agerbæk, M.O., Eriksen, E.F., Kragstrup, J., Mosekilde, Le. & Melsen, F. (1991) A reconstruction of the remodelling cycle in normal human cortical iliac bone. Bone and Mineral, 12, 101–112. Available from: 10.1016/0169-6009(91)90039-3

4. Allen, P.J., Baumgartner, W., Brinkman, E., DeVries, R.J., Stewart, H.A., Aboagye, D.L., et al. (2018) Fin healing and regeneration in sturgeon. Journal of Fish Biology, 93, 917–930. Available from: 10.1111/jfb.13794

5. Allen, P.J., Hobbs, J.A., Cech, J.J., Jr., Van Eenennaam, J.P. & Doroshov, S.I. (2009) Using Trace Elements in Pectoral Fin Rays to Assess Life History Movements in Sturgeon: Estimating Age at Initial Seawater Entry in Klamath River Green Sturgeon. Transactions of the American Fisheries Society, 138, 240–250. Available from: 10.1577/T08-061.1

6. Almany Magal, R., Reznikov, N., Shahar, R. & Weiner, S. (2014) Three-dimensional structure of minipig fibrolamellar bone: Adaptation to axial loading. Journal of Structural Biology, 186, 253–264. Available from: 10.1016/j.jsb.2014.03.007

7. Apschner, A., Schulte-Merker, S. & Witten, P.E. (2011) Not All Bones are Created Equal – Using Zebrafish and Other Teleost Species in Osteogenesis Research. In: Methods in Cell Biology. Elsevier, pp. 239–255. Available from: 10.1016/B978-0-12-381320-6.00010-2

8. Ashique, A.M., Atake, O.J., Ovens, K., Guo, R., Pratt, I.V., Detrich, H.W., et al. (2022) Bone microstructure and bone mineral density are not systemically different in Antarctic icefishes and related Antarctic notothenioids. Journal of Anatomy, 240, 34–49. Available from: 10.1111/joa.13537

9. Atkins, A., Dean, M.N., Habegger, M.L., Motta, P.J., Ofer, L., Repp, F., et al. (2014) Remodeling in bone without osteocytes: Billfish challenge bone structure–function paradigms. Proceedings of the National Academy of Sciences, 111, 16047–16052. Available from: 10.1073/pnas.1412372111

10. Atkins, A., Reznikov, N., Ofer, L., Masic, A., Weiner, S. & Shahar, R. (2015) The three-dimensional structure of anosteocytic lamellated bone of fish. Acta Biomaterialia, 13, 311–323. Available from: 10.1016/j.actbio.2014.10.025

11. Bakhshalizadeh, S., Nasibulina, B., Kurochkina, T., Ali, A., Mora-Medina, R. & Ayala-Soldado, N. (2023a) Multivariate analysis of trace elements in starry sturgeon (*Acipenser stellatus*) spine in different areas of the Caspian Sea. Marine Pollution Bulletin, 194, 115289. Available from: 10.1016/j.marpolbul.2023.115289

12. Bakhshalizadeh, S., Nasibulina, B.M., Kurochkin, T.F., Ali, A.M. & Hermann, T.W. (2023b) Fin-spine microchemistry discriminates regional stocks of Caspian Sea starry sturgeon. Estuarine, Coastal and Shelf Science, 292, 108483. Available from: 10.1016/j.ecss.2023.108483

13. Barrera, J.W., Le Cabec, A. & Barak, M.M. (2016) The orthotropic elastic properties of fibrolamellar bone tissue in juvenile white tailed deer femora. Journal of Anatomy, 229, 568–576. Available from: 10.1111/joa.12500

14. Bemis, W.E., Findeis, E.K. & Grande, L. (1997) An overview of Acipenseriformes. Environmental Biology of Fishes, 48, 25–71. Available from: 10.1023/A:1007370213924

15. Benton, M., Donoghue, P., Vinther, J., Asher, R., Friedman, M. & Near, T. (2015) Constraints on the timescale of animal evolutionary history. Palaeontologia Electronica, 18. Available from: 10.26879/424

16. Bonfield, W. & Grynpas, M.D. (1977) Anisotropy of the Young’s modulus of bone. Nature, 270, 453–454. Available from: 10.1038/270453a0

17. Brownstein, C.D. & Near, T.J. (2025) Toward a Phylogenetic Taxonomy of Sturgeons (Acipenseriformes: Acipenseridae). Bulletin of the Peabody Museum of Natural History, 66, 3–23. Available from: 10.3374/014.066.0101

18. Bruch, R.M., Campana, S.E., Davis Foust, S.L., Hansen, M.J. & Janssen, J. (2009) Lake Sturgeon Age Validation using Bomb Radiocarbon and Known Age Fish. Transactions of the American Fisheries Society, 138, 361–372. Available from: 10.1577/T08-098.1

19. de Buffrénil, V., Clarac, F., Fau, M., Martin, S., Martin, B., Pellé, E., et al. (2015) Differentiation and growth of bone ornamentation in vertebrates: A comparative histological study among the Crocodylomorpha. Journal of Morphology, 276, 425–445. Available from: 10.1002/jmor.20351

20. Carter, D.R., Mikić, B. & Padian, K. (1998) Epigenetic mechanical factors in the evolution of long bone epiphyses. Zoological Journal of the Linnean Society, 123, 163–178. Available from: 10.1111/j.1096-3642.1998.tb01298.x

21. Cattaneo, C., DiMartino, S., Scali, S., Craig, O.E., Grandi, M. & Sokol, R.J. (1999) Determining the human origin of fragments of burnt bone: a comparative study of histological, immunological and DNA techniques. Forensic Science International, 102, 181–191. Available from: 10.1016/S0379-0738(99)00059-6

22. Clarac, F., Goussard, F., Teresi, L., Buffrénil, Vde. & Sansalone, V. (2017) Do the ornamented osteoderms influence the heat conduction through the skin? A finite element analysis in Crocodylomorpha. Journal of Thermal Biology, 69, 39–53. Available from: 10.1016/j.jtherbio.2017.06.003

23. Clifton, K.B., Yan, J., Mecholsky Jr., J.J. & Reep, R.L. (2008) Material properties of manatee rib bone. Journal of Zoology, 274, 150–159. Available from: 10.1111/j.1469-7998.2007.00366.x

24. Cohen, J. & Harris, W.H. (1958) The Three-Dimensional Anatomy of Haversian Systems. JBJS, 40, 419.

25. Cohen, L., Dean, M., Shipov, A., Atkins, A., Monsonego-Ornan, E. & Shahar, R. (2012) Comparison of structural, architectural and mechanical aspects of cellular and acellular bone in two teleost fish. Journal of Experimental Biology, 215, 1983–1993. Available from: 10.1242/jeb.064790

26. Congiu, T. & Pazzaglia, U.E. (2011) The Sealed Osteons of Cortical Diaphyseal Bone. Early Observations Revisited With Scanning Electron Microscopy. The Anatomical Record, 294, 193–198. Available from: 10.1002/ar.21309

27. Cooper, D.M.L., Matyas, J.R., Katzenberg, M.A. & Hallgrimsson, B. (2004) Comparison of Microcomputed Tomographic and Microradiographic Measurements of Cortical Bone Porosity. Calcified Tissue International, 74, 437–447. Available from: 10.1007/s00223-003-0071-z

28. Currey, J. (2008) Collagen and the Mechanical Properties of Bone and Calcified Cartilage. In: Fratzl, P. (Ed.) Collagen: Structure and Mechanics. Boston, MA: Springer US, pp. 397–420. Available from: 10.1007/978-0-387-73906-9_14

29. Currey, J.D. (1999) What determines the bending strength of compact bone? Journal of Experimental Biology, 202, 2495–2503. Available from: 10.1242/jeb.202.18.2495

30. Currey, J.D. (2002) Bones: Structure and Mechanics. Princeton University Press

31. Currey, J.D. & Shahar, R. (2013) Cavities in the compact bone in tetrapods and fish and their effect on mechanical properties. Journal of Structural Biology, 183, 107–122. Available from: 10.1016/j.jsb.2013.04.012

32. Currey, J.D., Zioupos, P., Peter, D. & Casinos, A. (2001) Mechanical properties of nacre and highly mineralized bone. Proceedings of the Royal Society B: Biological Sciences, 268, 107–111. Available from: 10.1098/rspb.2000.1337

33. Davesne, D., Meunier, F.J., Friedman, M., Benson, R.B.J. & Otero, O. (2018) Histology of the endothermic opah ( *Lampris* sp.) suggests a new structure–function relationship in teleost fish bone. Biology Letters, 14, 20180270. Available from: 10.1098/rsbl.2018.0270

34. Davesne, D., Schmitt, A.D., Fernandez, V., Benson, R.B.J. & Sanchez, S. (2020) Three dimensional characterization of osteocyte volumes at multiple scales, and its relationship with bone biology and genome evolution in ray finned fishes. Journal of Evolutionary Biology, 33, 808–830. Available from: 10.1111/jeb.13612

35. Dean, M.N. & Shahar, R. (2012) The structure-mechanics relationship and the response to load of the acellular bone of neoteleost fish: a review: Acellular bone structure-mechanics. Journal of Applied Ichthyology, 28, 320–329. Available from: 10.1111/j.1439-0426.2012.01991.x

36. van Der Meulen, M.C.H., Beaupré, G.S. & Carter, D.R. (1993) Mechanobiologic influences in long bone cross-sectional growth. Bone, 14, 635–642. Available from: 10.1016/8756-3282(93)90085-O

37. Dillman, C.B. & Hilton, E.J. (2014) Anatomy and early development of the pectoral girdle, fin, and fin spine of sturgeons (Actinopterygii: Acipenseridae). Journal of Morphology, 276, 241–260. Available from: 10.1002/jmor.20328

38. Donoghue, P.C.J. & Sansom, I.J. (2002) Origin and early evolution of vertebrate skeletonization. Microscopy Research and Technique, 59, 352–372. Available from: 10.1002/jemt.10217

39. Doube, M. (2022) Closing cones create conical lamellae in secondary osteonal bone. Royal Society Open Science, 9, 220712. Available from: 10.1098/rsos.220712

40. Du, K., Stöck, M., Kneitz, S., Klopp, C., Woltering, J.M., Adolfi, M.C., et al. (2020) The sterlet sturgeon genome sequence and the mechanisms of segmental rediploidization. Nature Ecology & Evolution, 4, 841–852. Available from: 10.1038/s41559-020-1166-x

41. Eastoe, J.E. & Eastoe, B. (1954) The organic constituents of mammalian compact bone. Biochemical Journal, 57, 453–459. Available from: 10.1042/bj0570453

42. Enlow, D.H. (1962) A Study of the Post-Natal Growth and Remodeling of Bone. American Journal of Anatomy, 110, 79–101. Available from: 10.1002/aja.1001100202

43. Enlow, D.H. (1966) An Evaluation of the Use of Bone Histology in Forensic Medicine and Anthropology. In: Evans, F.G. (Ed.) Studies on the Anatomy and Function of Bone and Joints. Berlin, Heidelberg: Springer, pp. 93–112. Available from: 10.1007/978-3-642-99909-3_7

44. Estefa, J., Tafforeau, P., Clement, A.M., Klembara, J., Niedźwiedzki, G., Berruyer, C., et al. (2021) New light shed on the early evolution of limb-bone growth plate and bone marrow. eLife, 10, e51581. Available from: 10.7554/eLife.51581

45. Ferrón, H.G., Ballell, A., Botella, H. & Martínez-Pérez, C. (2021) Biomechanics of Machaeracanthus pectoral fin spines provide evidence for distinctive spine function and lifestyle among early chondrichthyans. Journal of Vertebrate Paleontology, 41, e2090260. Available from: 10.1080/02724634.2021.2090260

46. Findeis, E.K. (1993) *Skeletal anatomy of the North American shovelnose sturgeon Scaphirhynchus platorynchus (Rafinesque 1820) with comparisons to other Acipenseriformes*. Ph.D. University of Massachusetts Amherst. Available from: https://www.proquest.com/docview/304052521/abstract/E99E223E7884645PQ/1. Accessed 10 October 2025

47. Findeis, E.K. (1997) Osteology and phylogenetic interrelationships of sturgeons (Acipenseridae). Environmental Biology of Fishes, 48, 73–126. Available from: 10.1023/A:1007372832213

48. Fletcher, J.W.A., Williams, S., Whitehouse, M.R., Gill, H.S. & Preatoni, E. (2018) Juvenile bovine bone is an appropriate surrogate for normal and reduced density human bone in biomechanical testing: a validation study. Scientific Reports, 8, 10181. Available from: 10.1038/s41598-018-28155-w

49. Francillon-Vieillot, H., De Buffrénil, V., Castanet, J., Géraudie, J., Meunier, F.J., Sire, J.Y., et al. (1990) Microstructure and Mineralization of Vertebrate Skeletal Tissues. In: Carter, J.G. (Ed.) Skeletal Biomineralization: Patterns, Processes and Evolutionary Trends. Washington, D. C.: American Geophysical Union, pp. 175–234. Available from: 10.1029/SC005p0175

50. Galea, G.L., Zein, M.R., Allen, S. & Francis-West, P. (2021) Making and shaping endochondral and intramembranous bones. Developmental Dynamics, 250, 414–449. Available from: 10.1002/dvdy.278

51. Haines, R.W. (1942) The Evolution of Epiphyses and of Endochondral Bone. Biological Reviews, 17, 267–292. Available from: 10.1111/j.1469-185X.1942.tb00440.x

52. Hall, B.K. (2015) Bones and Cartilage: Developmental and Evolutionary Skeletal Biology. 2nd edition. Elsevier. Available from: 10.1016/C2013-0-00143-0

53. Havers, C. (1691) Osteologia nova, or, Some new observations of the bones, and the parts belonging to them, with the manner of their accretion, and nutrition, communicated to the Royal Society in several discourses to which is added a fifth discourse of the cartilages / by Clopton Havers *…* Available from: http://name.umdl.umich.edu/B24042.0001.001

54. Heřt, J., Fiala, P. & Petrtýl, M. (1994) Osteon orientation of the diaphysis of the long bones in man. Bone, 15, 269–277. Available from: 10.1016/8756-3282(94)90288-7

55. Hill, R.V. (2006) Comparative anatomy and histology of xenarthran osteoderms. Journal of Morphology, 267, 1441–1460. Available from: 10.1002/jmor.10490

56. Hilton, E.J., Grande, L. & Bemis, W.E. (2011) Skeletal Anatomy of the Shortnose Sturgeon, Acipenser brevirostrum Lesueur, 1818, and the Systematics of Sturgeons (Acipenseriformes, Acipenseridae). *Fieldiana Life and Earth Sciences*, 2011, 1–168. Available from: 10.3158/2158-5520-3.1.1

57. Hilton, E.J., Grande, L. & Jin, F. (2021) Redescription of † Yanosteus longidorsalis Jin et al., 1995 (Chondrostei, Acipenseriformes, †Peipiaosteidae) from the Early Cretaceous of China. Journal of Paleontology, 95, 170–183. Available from: 10.1017/jpa.2020.80

58. Hirasawa, T. & Kuratani, S. (2015) Evolution of the vertebrate skeleton: morphology, embryology, and development. Zoological Letters, 1, 2. Available from: 10.1186/s40851-014-0007-7

59. Höch, R., Schneider, R.F., Kickuth, A., Meyer, A. & Woltering, J.M. (2021) Spiny and soft-rayed fin domains in acanthomorph fish are established through a BMP-gremlin-shh signaling network. Proceedings of the National Academy of Sciences, 118, e2101783118. Available from: 10.1073/pnas.2101783118

60. Horton, J.M. & Summers, A.P. (2009) The material properties of acellular bone in a teleost fish. Journal of Experimental Biology, 212, 1413–1420. Available from: 10.1242/jeb.020636

61. Hughes, D.R., Bassett, J.R. & Moffat, L.A. (1994) Histological identification of osteocytes in the allegedly acellular bone of the sea breams Acanthopagrus australis, Pagrus auratus and Rhabdosargus sarba (Sparidae, Perciformes, Teleostei). Anatomy and Embryology, 190, 163–179. Available from: 10.1007/BF00193413

62. Hung, S.S.O. (2017) Recent advances in sturgeon nutrition. Animal Nutrition, 3, 191–204. Available from: 10.1016/j.aninu.2017.05.005

63. Huysseune, A. & Sire, J.-Y. (1998) Evolution of patterns and processes in teeth and tooth-related tissues in non-mammalian vertebrates. European Journal of Oral Sciences, 106, 437–481. Available from: 10.1111/j.1600-0722.1998.tb02211.x

64. Infrared Spectra Library | The Helen and Martin Kimmel Center (2021). Available from: https://centers.weizmann.ac.il/kimmel-arch/infrared-spectra-library. Accessed 22 March 2026

65. Janis, C.M., Devlin, K., Warren, D.E. & Witzmann, F. (2012) Dermal bone in early tetrapods: a palaeophysiological hypothesis of adaptation for terrestrial acidosis. Proceedings of the Royal Society B: Biological Sciences, 279, 3035–3040. Available from: 10.1098/rspb.2012.0558

66. Jerve, A., Bremer, O., Sanchez, S. & Ahlberg, P. (2017) Morphology and histology of acanthodian fin spines from the late Silurian Ramsåsa E locality, Skåne, Swede. Palaeontologia Electronica [Preprint]. Available from: 10.26879/749

67. Jerve, A., Qu, Q., Sanchez, S., Blom, H. & Ahlberg, P.E. (2016) Three-dimensional paleohistology of the scale and median fin spine of Lophosteus superbus (Pander 1856). PeerJ, 4, e2521. Available from: 10.7717/peerj.2521

68. Johanson, Z., Liston, J., Davesne, D., Challands, T. & Meredith Smith, M. (2022) Mechanisms of dermal bone repair after predatory attack in the giant stem-group teleost Leedsichthys problematicus Woodward, 1889a (Pachycormiformes). Journal of Anatomy, 241, 393–406. Available from: 10.1111/joa.13689

70. Kaatz, I.M., Stewart, D.J., Rice, A.N. & Lobel, P.S. (2010) Differences in pectoral fin spine morphology between vocal and silent clades of catfishes (Order Siluriformes): Ecomorphological implications. Current Zoology, 56, 73–89. Available from: 10.1093/czoolo/56.1.73

71. Kalish-Achrai, N., Monsonego-Ornan, E. & Shahar, R. (2017) Structure, composition, mechanics and growth of spines of the dorsal fin of blue tilapia Oreochromis aureus and common carp Cyprinus carpio. Journal of Fish Biology, 90, 2073–2096. Available from: 10.1111/jfb.13287

72. Kim, J.-N., Lee, J.-Y., Shin, K.-J., Gil, Y.-C., Koh, K.-S. & Song, W.-C. (2015) Haversian system of compact bone and comparison between endosteal and periosteal sides using three-dimensional reconstruction in rat. Anatomy & Cell Biology, 48, 258–261. Available from: 10.5115/acb.2015.48.4.258

73. Kornblum, S.S. & Kelly, P.J. (1964) The Lacunae and Haversian Canals in Tibial Cortical Bone from Ischemic and Non-Ischemic Limbs: A COMPARATIVE MICRORADIOGRAPHIC STUDY. JBJS, 46, 797.

74. Kronenberg, H.M. (2003) Developmental regulation of the growth plate. Nature, 423, 332–336. Available from: 10.1038/nature01657

75. Kubicek, K.M., Britz, R. & Conway, K.W. (2025) Heterochrony leads to evolutionary novelty: the catfish pectoral-fin spine (Teleostei: Siluriformes). Biology Letters, 21, 20250038. Available from: 10.1098/rsbl.2025.0038

76. Locke, M. (2004) Structure of long bones in mammals. Journal of Morphology, 262, 546–565. Available from: 10.1002/jmor.10282

77. Lowenstam, H.A. & Weiner, S. (1989) On Biomineralization. Oxford University Press. Available from: 10.1093/oso/9780195049770.001.0001

78. Lundberg, J.G. & Aguilera, O. (2003) The late Miocene Phractocephalus catfish (Siluriformes: Pimelodidae) from Urumaco, Venezuela: additional specimens and reinterpretation as a distinct species. Neotropical Ichthyology, 1, 97–109. Available from: 10.1590/S1679-62252003000200004

79. Martens, M., van Audekercke, R., de Meester, P. & Mulier, J.C. (1980) The mechanical characteristics of the long bones of the lower extremity in torsional loading. Journal of Biomechanics, 13, 667–676. Available from: 10.1016/0021-9290(80)90353-X

80. Martin, R.B. & Burr, D.B. (1989) Structure, function, and adaptation of compact bone. New York: Raven Press

81. Meunier, F.J. & Gayet, M. (2020) Comparative morphology of the finlet spines of the extant Polypteridae (Osteichthyes; Cladistia). Systematic interest. Available from: 10.26028/CYBIUM/2020-441-003

82. Milgram, J., Rehav, K., Ibrahim, J., Shahar, R. & Weiner, S. (2023) The 3D organization of the mineralized scales of the sturgeon has structures reminiscent of dentin and bone: A FIB-SEM study. Journal of Structural Biology, 215, 108045. Available from: 10.1016/j.jsb.2023.108045

83. Morgan, E.F., Unnikrisnan, G.U. & Hussein, A.I. (2018) Bone Mechanical Properties in Healthy and Diseased States. Annual Review of Biomedical Engineering, 20, 119–143. Available from: 10.1146/annurev-bioeng-062117-121139

84. Mosca, K.C., Savoy, T.F., Roberts, J.B., Ingram, E.C., Schultz, E.T. & Baumann, H. (2025) Age structure and seasonal movement of Atlantic sturgeon (Acipenser oxyrinchus) aggregating in eastern Long Island Sound and the Connecticut River. Fishery Bulletin, 123, 127–142. Available from: 10.7755/FB.123.2.5

85. Moss, M.L. (1968) Comparative anatomy of vertebrate dermal bone and teeth: I. The epidermal co-participation hypothesis. Acta Anatomica, 71, 178–208. Available from: 10.1159/000143185

86. Nakano, T. (2015) Bone Tissue and Biomaterial Design Based on the Anisotropic Microstructure. In: Niinomi, M., Narushima, T., & Nakai, M. (Eds) Advances in Metallic Biomaterials. Berlin, Heidelberg: Springer Berlin Heidelberg (Springer Series in Biomaterials Science and Engineering), pp. 3–30. Available from: 10.1007/978-3-662-46836-4_1

87. Neary, J.J., Pracheil, B.M., Gabitov, R.I., Li, M.H. & Allen, P.J. (2024) The influence of water, diet, and temperature on 87Sr/86Sr in fin spines of juvenile Atlantic Sturgeon *Acipenser oxyrinchus oxyrinchus*. Journal of Experimental Marine Biology and Ecology, 570, 151973. Available from: 10.1016/j.jembe.2023.151973

88. Oliver, W.C. & Pharr, G.M. (1992) An improved technique for determining hardness and elastic modulus using load and displacement sensing indentation experiments. Journal of Materials Research, 7, 1564–1583. Available from: 10.1557/JMR.1992.1564

89. Patterson, C., Andrews, S.M., Miles, R.S. & Walker, A.D. (1979) Cartilage bones, dermal bones and membrane bones, or the exoskeleton versus the endoskeleton. In: Problems in Vertebrate Evolution: Essays Presented to Professor T. S. Westoll. London: Academic press for the Linnean society of London (Linnean society symposium series, 4)

90. Peng, Z., Ludwig, A., Wang, D., Diogo, R., Wei, Q. & He, S. (2007) Age and biogeography of major clades in sturgeons and paddlefishes (Pisces: Acipenseriformes). Molecular Phylogenetics and Evolution, 42, 854–862. Available from: 10.1016/j.ympev.2006.09.008

91. Price, S.A., Friedman, S.T. & Wainwright, P.C. (2015) How predation shaped fish: the impact of fin spines on body form evolution across teleosts. Proceedings of the Royal Society B: Biological Sciences, 282, 20151428. Available from: 10.1098/rspb.2015.1428

92. Prondvai, E., Stein, K.H.W., de Ricqlès, A. & Cubo, J. (2014) Development-based revision of bone tissue classification: the importance of semantics for science. Biological Journal of the Linnean Society, 112, 799–816. Available from: 10.1111/bij.12323

93. Qu, Q., Zhu, M. & Wang, W. (2013) Scales and Dermal Skeletal Histology of an Early Bony Fish Psarolepis romeri and Their Bearing on the Evolution of Rhombic Scales and Hard Tissues. PLoS ONE. Edited by V. Laudet, 8, e61485. Available from: 10.1371/journal.pone.0061485

94. Quan, H., Yang, W., Tang, Z., Ritchie, R.O. & Meyers, M.A. (2020) Active defense mechanisms of thorny catfish. Materials Today, 38, 35–48. Available from: 10.1016/j.mattod.2020.04.028

95. Raguin, E., Rechav, K., Brumfeld, V., Shahar, R. & Weiner, S. (2020) Unique three-dimensional structure of a fish pharyngeal jaw subjected to unusually high mechanical loads. Journal of Structural Biology, 211, 107530. Available from: 10.1016/j.jsb.2020.107530

96. Reznikov, N., Shahar, R. & Weiner, S. (2014) Bone hierarchical structure in three dimensions. Acta Biomaterialia, 10, 3815–3826. Available from: 10.1016/j.actbio.2014.05.024

97. Rho, J.-Y., Kuhn-Spearing, L. & Zioupos, P. (1998) Mechanical properties and the hierarchical structure of bone. Medical Engineering & Physics, 20, 92–102. Available from: 10.1016/S1350-4533(98)00007-1

98. de Ricqlès, A. (1975) Recherches paleohistologiques sur les os longs des tetrapodes. Annales de Paleontologie, 61, 51–129.

99. Saha, S. (1973) Anisotropic analysis of bone — Some two-dimensional problems. Journal of Biomechanics, 6, 641–650. Available from: 10.1016/0021-9290(73)90020-1

100. Scheyer, T.M., Schmid, L., Furrer, H. & Sánchez-Villagra, M.R. (2014) An assessment of age determination in fossil fish: the case of the opercula in the Mesozoic actinopterygian Saurichthys. Swiss Journal of Palaeontology, 133, 243–257. Available from: 10.1007/s13358-014-0068-4

101. Seidel, M.R. (1979) The Osteoderms of the American Alligator and Their Functional Significance. Herpetologica, 35, 375–380.

102. Seto, J., Gupta, H.S., Zaslansky, P., Wagner, H.D. & Fratzl, P. (2008) Tough Lessons From Bone: Extreme Mechanical Anisotropy at the Mesoscale. Advanced Functional Materials, 18, 1905–1911. Available from: 10.1002/adfm.200800214

103. Singh, I.J., Tonna, E.A. & Gandel, C.P. (1974) A comparative histological study of mammalian bone. Journal of Morphology, 144, 421–437. Available from: 10.1002/jmor.1051440404

104. Sire, J., Donoghue, P.C.J. & Vickaryous, M.K. (2009) Origin and evolution of the integumentary skeleton in non tetrapod vertebrates. Journal of Anatomy, 214, 409–440. Available from: 10.1111/j.1469-7580.2009.01046.x

105. Sire, J. & Huysseune, A. (2003) Formation of dermal skeletal and dental tissues in fish: a comparative and evolutionary approach. Biological Reviews, 78, 219–249. Available from: 10.1017/S1464793102006073

106. Skedros, J.G., Clark, G.C., Sorenson, S.M., Taylor, K.W. & Qiu, S. (2011) Analysis of the Effect of Osteon Diameter on the Potential Relationship of Osteocyte Lacuna Density and Osteon Wall Thickness. The Anatomical Record, 294, 1472–1485. Available from: 10.1002/ar.21452

107. Skedros, J.G., Henrie, T.R., Doutré, M.S. & Bloebaum, R.D. (2018) Sealed osteons in animals and humans: low prevalence and lack of relationship with age. Journal of Anatomy, 232, 824–835. Available from: 10.1111/joa.12786

108. Smith-Vaniz, W.F., Kaufman, L.S. & Glowacki, J. (1995) Species-specific patterns of hyperostosis in marine teleost fishes. Marine Biology, 121, 573–580. Available from: 10.1007/BF00349291

109. Talts, J.F., Pfeifer, A., Hofmann, F., Hunziker, E.B., Zhou, X.-H., Aszódi, A., et al. (1998) Endochondral Ossification Is Dependent on the Mechanical Properties of Cartilage Tissue and on Intracellular Signals in Chondrocytes. Annals of the New York Academy of Sciences, 857, 74–85. Available from: 10.1111/j.1749-6632.1998.tb10108.x

110. Tappen, N.C. (1977) Three-dimensional studies of resorption spaces and developing osteons. American Journal of Anatomy, 149, 301–331. Available from: 10.1002/aja.1001490302

111. Thacker, C.E. & Near, T.J. (2025) Phylogeny, biology, and evolution of acanthopterygian fish clades. Reviews in Fish Biology and Fisheries, 35, 805–845. Available from: 10.1007/s11160-025-09935-w

112. Tomes, J. & Morgan, C.G.D. (1853) IV. Observations on the structure and development of bone. Philosophical Transactions of the Royal Society of London, 109–139. Available from: 10.1098/rstl.1853.0004

113. Toppe, J., Albrektsen, S., Hope, B. & Aksnes, A. (2007) Chemical composition, mineral content and amino acid and lipid profiles in bones from various fish species. Comparative Biochemistry and Physiology Part B: Biochemistry and Molecular Biology, 146, 395–401. Available from: 10.1016/j.cbpb.2006.11.020

114. Vanscoy, T., Lundberg, J.G. & Luckenbill, K.R. (2015) Bony ornamentation of the catfish pectoral-fin spine: comparative and developmental anatomy, with an example of fin-spine diversity using the Tribe Brachyplatystomini (Siluriformes, Pimelodidae). Proceedings of the Academy of Natural Sciences of Philadelphia, 164, 177–212. Available from: 10.1635/053.164.0107

115. Vickaryous, M.K. & Hall, B.K. (2006) Osteoderm morphology and development in the nine-banded armadillo, Dasypus novemcinctus (Mammalia, Xenarthra, Cingulata). Journal of Morphology, 267, 1273–1283. Available from: 10.1002/jmor.10475

116. Wan, Q.-Q., Qin, W.-P., Ma, Y.-X., Shen, M.-J., Li, J., Zhang, Z.-B., et al. (2021) Crosstalk between Bone and Nerves within Bone. Advanced Science, 8, 2003390. Available from: 10.1002/advs.202003390

117. Watson, D.M.S. (1937) II - The Acanthodian fishes. *Philosophical Transactions of the Royal Society of London. B*, Biological Sciences, 228, 49–146. Available from: 10.1098/rstb.1937.0009

118. Weigele, J. & Franz Odendaal, T.A. (2016) Functional bone histology of zebrafish reveals two types of endochondral ossification, different types of osteoblast clusters and a new bone type. Journal of Anatomy, 229, 92–103. Available from: 10.1111/joa.12480

119. Weiner, S. & Wagner, H.D. (1998) THE MATERIAL BONE: Structure-Mechanical Function Relations. Annual Review of Materials Research, 28, 271–298. Available from: 10.1146/annurev.matsci.28.1.271

120. Wilga, C.D. & Lauder, G.V. (1999) Locomotion in sturgeon: function of the pectoral fins. Journal of Experimental Biology, 202, 2413–2432. Available from: 10.1242/jeb.202.18.2413

121. Witten, P.E. & Hall, B.K. (2002) Differentiation and growth of kype skeletal tissues in anadromous male Atlantic salmon (Salmo salar). The International Journal of Developmental Biology, 46, 719–730.

122. Witzmann, F., Scholz, H., Müller, J. & Kardjilov, N. (2010) Sculpture and vascularization of dermal bones, and the implications for the physiology of basal tetrapods. Zoological Journal of the Linnean Society, 160, 302–340. Available from: 10.1111/j.1096-3642.2009.00599.x

123. Witzmann, F. & Soler-Gijón, R. (2010) The bone histology of osteoderms in temnospondyl amphibians and in the chroniosuchian Bystrowiella. Acta Zoologica, 91, 96–114. Available from: 10.1111/j.1463-6395.2008.00385.x

124. Zedda, M. & Babosova, R. (2021) Does the osteon morphology depend on the body mass? A scaling study on macroscopic and histomorphometric differences between cow (Bos taurus) and sheep (Ovis aries). Zoomorphology, 140, 169–181. Available from: 10.1007/s00435-021-00516-6

