## Supplementary materials for "Functional Adaptations for Load-Bearing in a Dermal Bone: The Pectoral Fin Spine of the Russian Sturgeon (*Huso gueldenstaedtii*)"

### Supplementary material

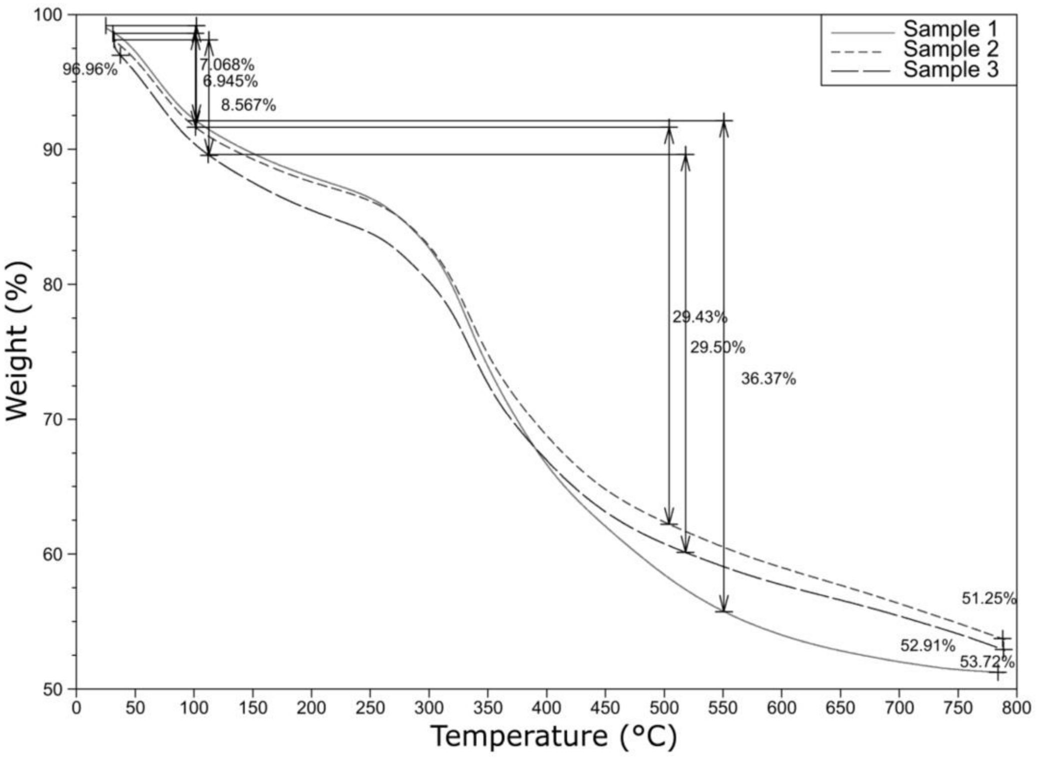


**Supplementary material 1.** Thermogravimetric analysis (TGA) of the PFS. The graph shows the curves of the heated PFS bone as loss of material happens.

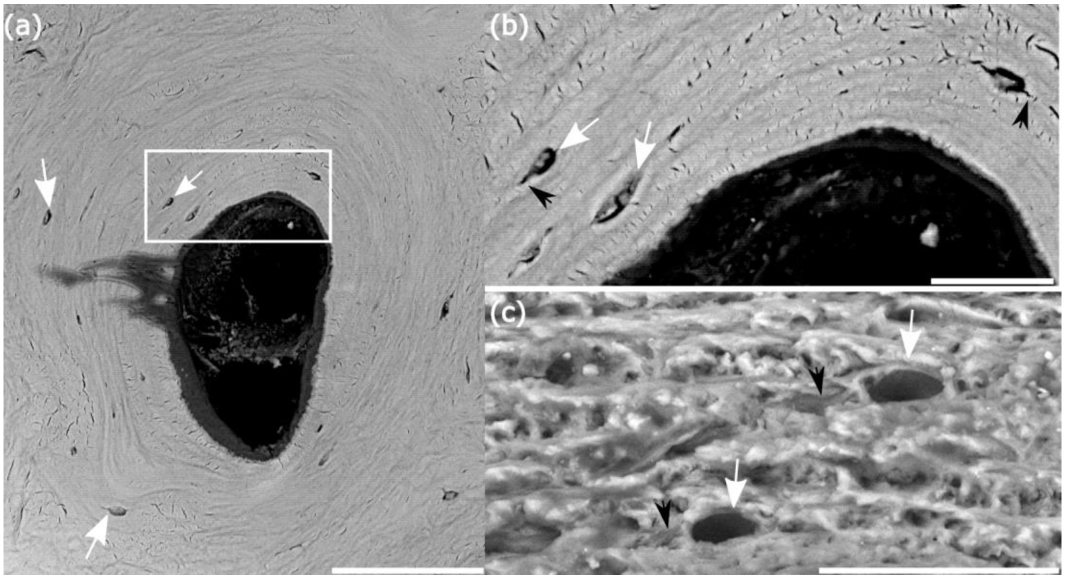


**Supplementary material 2**. Backscatter electron detector images of a polished section of the central region (a and b) and a fracture surface of the peripheral region (c) showing the presence of osteocytic lacunae. All surfaces were prepared in transverse plane. Voids with diameters consistent with osteocytic lacunae are shown in all images (white arrows). The location of osteocytic lacunae between the layers of bone forming the dermal osteonal bone are seen in (a), and canaliculi (black arrow) can be seen extending from the osteocytic lacunae in the region of the white square, which is shown at higher magnification in (b). Osteocytic lacunae (white arrows) and canaliculi (black arrow) are also seen in the peripheral region (c). Scale bars: a) 100µm; b) 25µm; c) 30µm.
